# A polarized nuclear position is required for correct division plane specification during maize stomatal development

**DOI:** 10.1101/2022.08.26.505454

**Authors:** M. Arif Ashraf, Le Liu, Michelle R. Facette

## Abstract

Asymmetric cell division generates new cell types and is a feature of development in multicellular organisms. Prior to asymmetric cell division, cell polarity is established. *Zea mays* stomatal development serves as an excellent plant model system for asymmetric cell division, especially the asymmetric division of the subsidiary mother cell (SMC). In SMCs, the nucleus migrates to a polar location after the accumulation of polarly localized proteins, but before the appearance of the preprophase band. We examined a mutant of the outer nuclear membrane protein, which is part of the LINC (linker of nucleoskeleton and cytoskeleton) complex that localizes to the nuclear envelope in interphase cells. Previously, *mlks2* (*maize linc kash sine-like2*) was observed to have abnormal stomata. We confirmed and identified the precise defects that lead to abnormal asymmetric divisions. Proteins that are polarly localized in SMCs prior to division polarize normally in *mlks2*. However, polar localization of the nucleus is sometimes impaired, even in cells that have otherwise normal polarity. This leads to a misplaced preprophase band and atypical division planes. MLKS2 is localized to mitotic structures, however the structure of the preprophase band, spindle and phragmoplast appeared normal in *mlks2*. Timelapse imaging revealed that *mlks2* has defects in pre-mitotic nuclear migration towards the polarized site, and unstable position at the division site after formation of the preprophase band. We show that nuclear envelope proteins promote pre-mitotic nuclear migration and stable nuclear position, and that the position of the nucleus influences division plane establishment in asymmetrically dividing cells.

**One-sentence summary:** Nuclear movement and positioning prior to asymmetric cell division is required for deciding the future division site.

## Introduction

Nuclear membrane proteins reside in either the inner or outer nuclear membrane. Certain nuclear membrane proteins help to maintain the connection between nuclei and the rest of the cell. Because of the nature of this specific class of protein, they are known as LINC (<u>L</u>inker of <u>n</u>ucleoskeleton and <u>c</u>ytoskeleton) complexes. The LINC complex consists of KASH (<u>K</u>larsicht, <u>A</u>NC-1, and <u>S</u>YNE <u>h</u>omology) domain containing proteins in the outer membrane, and SUN (<u>S</u>ad-1/<u>UN</u>C-84) domain containing proteins in the inner membrane (Tapley and Starr, 2013; Gumber et al., 2019a; Groves et al., 2020). KASH and SUN domain-containing proteins interact with each other in the perinuclear space. KASH domain-containing proteins interact with cytoskeleton and SUN domain-containing proteins are associated with nuclear lamins and chromatin (Supplemental Figure 1A). As a result, the LINC complexes relay signals and act as a vehicle for nuclear mechanosensing (Uzer et al., 2016; Fal et al., 2017). At the same time, interaction of the nucleus with the cytoskeleton through the LINC complex facilitates nuclear movement (Tamura et al., 2013; Starr, 2019). In plants, nuclear movement is an important event during pollen and root hair growth, and during asymmetric cell divisions that occur during the formation of the zygote, lateral roots and stomata (Gallagher and Smith, 2000; Kawashima et al., 2014; Kimata et al., 2016; Barro et al., 2019; Ashraf and Facette, 2020; Muroyama et al., 2020; Yi and Goshima, 2020; Brueggeman et al., 2022). In the aforementioned cells, the nucleus takes on a polarized position, where its asymmetrical localization is a key part of the growth or division process.

During symmetric and asymmetric cell divisions, the cortical division zone, which later narrows to the cortical division site, is specified (Smertenko et al., 2017). The preprophase band (PPB) marks the cortical division zone at the G2/M transition, followed by other markers such as TRMs, TAN1, POKs, KCBP, RAN-GAP and MAP65-4 (Rasmussen and Bellinger, 2018). Prior to the appearance of the PPB, the nucleus moves to the future division site. Symmetric divisions tend to follow geometrical rules, with the nucleus centered prior to division (Martinez et al., 2018; Moukhtar et al., 2019). Nuclear movement to the division site is especially apparent in asymmetric divisions. Nuclear movement prior to asymmetric cell division is stimulated by an intrinsic or extrinsic polarizing cue, and often results in the polarized accumulation of proteins or organelles (Li, 2013). In some cases, asymmetrically dividing cells produce two volumetrically equal daughter cells and nuclear movement is not apparent (Facette et al., 2019). In contrast, when an asymmetrically dividing cell yields two volumetrically distinct daughter cells, nuclear movement is visible prior to mitosis. Examples of volumetrically distinct plant cell divisions include the first asymmetric cell division of the zygote, protonemal branch formation, and stomatal divisions in grass and non-grass species (Gallagher and Smith, 2000; Kimata et al., 2016; Ashraf and Facette, 2020; Yi and Goshima, 2020). Nuclear movement to the polarized site can be actin or microtubule dependent (Kimata et al., 2016; Barro et al., 2019; Ashraf and Facette, 2020; Muroyama et al., 2020; Yi and Goshima, 2020). Movement of the nucleus happens prior to establishment of the cortical division site. This implies that nuclear position may feed into determining the future site of division. It has been shown that cells that lack a preprophase band (through genetic perturbation), can still correctly specify the cortical division zone, and while the newly formed cell wall may have a more variable angle, they are essentially correct (Schaefer et al., 2017). It has been suggested that the preprophase band may stabilize or reinforce the division site, rather than specify it (Livanos and Müller, 2019). This implies that other cues feed into establishment of the cortical division site - but what are they? This is especially relevant in volumetrically distinct asymmetric cell divisions, where cues such as cell geometry may conflict with polarity cues.

Genetic perturbations that alter cellular polarity result in mislocalized nuclei, misaligned PPBs, and aberrant division planes during asymmetric divisions in several cell types (Gallagher and Smith, 2000; Kimata et al., 2019). Physical displacement of the nucleus via centrifugal force can also result in misaligned PPBs (Murata and Wada, 1991). These observations suggest that nuclear position can influence PPB placement. But, in these studies, cellular manipulation took place upstream (cell polarity establishment, organelle morphology, external cues) of the nuclear movement and PPB formation. For this reason, it is difficult to narrow down whether nuclear displacement and PPB establishment are a consequence of a cell polarity defect, or alteration of organelle morphology or an external physical cue? At this point, the role of nuclear position in PPB placement is still a question to answer and it requires genetic and cell biology tools to manipulate the nuclear movement exclusively without hampering upstream cellular events.

To tackle this fundamental question, we have targeted LINC complex components from the outer nuclear membrane protein, known to interact with the cytoskeleton, to eliminate the effect of upstream cellular events. In this study, we used a reverse genetics approach to investigate the function of an outer nuclear membrane protein, a component of the LINC complex, MLKS2 (<u>M</u>aize <u>L</u>INC <u>K</u>ASH At<u>S</u>INE-like), during asymmetric cell division in maize stomatal development. Interestingly, *mlks2* loss of function mutants have abnormally shaped subsidiary cells, products of asymmetric cell division during stomatal development (Gumber et al., 2019b). These mutants provided a tool that we used to dissect the role of outer nuclear membrane proteins during asymmetric cell division. Our cell biology studies show that defects in nuclear polarization in *mlks2* mutants lead to the formation of misplaced PPBs and subsequent mitotic structures (spindles, phragmoplasts, and cell plates). Timelapse imaging indicates that *mlks2* is defective in premitotic nuclear migration and maintenance of nuclear position after PPB establishment, however once the PPB is established division proceeds normally. Our work provides evidence that nuclear position plays a key role in PPB placement and subsequent division plane orientation.

## Results

### MLKS2 is required for asymmetric cell division during stomatal development

Previously, it was shown that *mlks2* mutants, mutations in an outer nuclear membrane proteins, have defects during meiosis, pollen viability, nuclear shape, and stomatal development (Gumber et al., 2019b). We reexamined two mutant lines, *mlks2-1* and *mlks2-2*, with Mu insertions at different locations in exon 1 (Supplemental Figure S1B), for stomatal phenotypes in expanded leaf 3 (Figure 1). Consistent with previous results (Gumber et al., 2019b), both mutant alleles result in aberrant subsidiary cells (Figure 1), extra interstomatal cells and aborted guard mother cells (Supplemental Figure S2). The variety of stomatal phenotypes in *mlks2* implies that abnormal divisions occur at multiple steps. For the correct orientation of the first asymmetric cell division that generates the guard mother cell (GMC), the nucleus polarizes shootward, followed by an asymmetric division generating a small GMC and an interstomatal cell (Supplemental Figure S3, S4A). Compared to wild type, mispositioned nuclei and preprophase bands are observed in *mlks2-1* during this initial division (Supplemental Figure S3A, B). The resulting misoriented division leads to incorrect fate specification of the GMC or an extra interstomatal cell (Supplemental Figure S3, S4). After GMCs are specified, SMCs are recruited from lateral pavement cells. The SMCs then divide asymmetrically to form a subsidiary cell and a pavement cell (Supplemental Figure S4B). Aberrant subsidiary cells, extra interstomatal cells, and aborted guard mother cells were the most common defects observed, and were quantified, however we also observed rare defects we never see in wild type. Examples of different abnormal stomatal phenotypes observed in *mlks2* mutants, and the aberrant divisions that result in such defects (Supplemental Figure S4).

**Figure 1.**
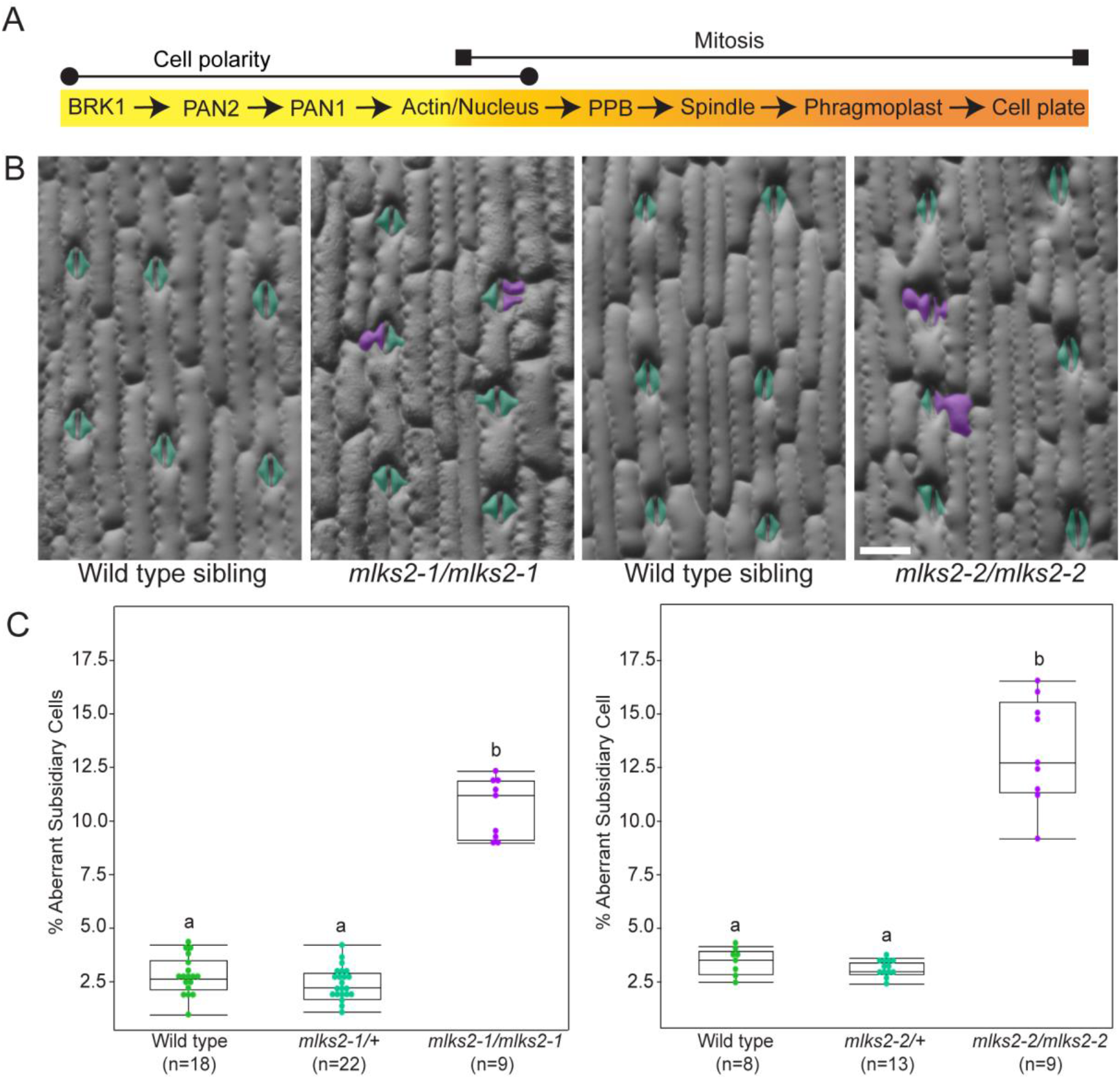
MLKS2 is required for asymmetric cell division. A, Timeline for cell polarity establishment and mitotic events during the formation of subsidiary cells. B, Subsidiary cell phenotypes of *mlks2-1* and *mlks2-2*. Cyan and magenta highlight indicate the normal and aberrant subsidiary cells, respectively. Scale bar = 0.1 mm. Same scale bar is applicable to all images. C, Quantification of aberrant subsidiary cells of *mlks2-1* and *mlks2-2*. Number of plants (n) used per genotypes are mentioned. At least 200 cells are observed from each individual plant. Statistical test is performed based on Tukey’s Honest Test. Groups labelled with the same letter are not statistically different from each other (alpha = 0.05).

The number of aberrant cells varies depending on growth conditions and leaf number. Polarity mutants such as *pan1, pan2* and *brk1* have more abnormal stomatal cells in early-emerging leaves (unpublished data), while a previously characterized nuclear pore complex protein, *aladin1*, had more abnormal stomatal cells in late-emerging leaves (Best et al., 2021). To determine if *mlks2* mutants behave more like polarity mutants vs *aladin1*, we analyzed stomatal cells in leaf 3, leaf 5, leaf 7 and leaf 10, where leaf 3 is the third leaf to emerge from the whorl (Supplemental Figure S5). There are more aberrant subsidiary cells in earlier-emerging leaves (leaf 3 and 5) than later emerging leaves (leaf 7 and 10). Moreover, the frequency of aberrant subsidiary cells is higher in fast-growing summer field plants (12-22% in *mlks2-1* and 15-27% in *mlks2-2*) than slow-growing winter greenhouse grown plants (8-12% in *mlks2-1* and 9-16% in *mlks2-2*) (Figure 1; and Supplemental Figure S5). This indicates that faster growing plants in the summer field conditions have a higher frequency of aberrant subsidiary cells.

In *A. thaliana*, the related protein, SINE2, is required for stomatal closure (Groves et al., 2020).To determine if MLKS2 is similarly required for stomatal closure, we used gas exchange to determine if *mlks2* plants were able to change stomatal conductance (i.e., close their stomata) in response to darkness using leaf 3 from greenhouse-grown plants. The mutant alleles responded similarly to wild type (Supplemental Figure S6), suggesting that MLKS2 is not required for stomatal closure under our tested conditions.

### MLKS2 localizes at the nuclear membrane and with the mitotic apparatus

Previously, it was shown that MLKS2 localizes to the nuclear membrane, and this localization is dependent on the C-terminal transmembrane and KASH domains (Gumber et al., 2019b). We repeated this expression analysis by transiently expressing MLKS2-mNeonGreen in tobacco leaf pavement cells and also found a nuclear envelope localization (Figure 7A). Interestingly, previous studies demonstrated that ectopic expression of AtSUN1 (inner nuclear membrane protein and part of the LINC complex) not only localizes at the nuclear membrane, but also co-localizes with mitotic structures after nuclear envelope breakdown (Graumann and Evans, 2011; Oda and Fukuda, 2011). Since *mlks2* mutants have a defect in asymmetric cell division, we thought it was important to determine if MLKS2 can also localize to mitotic structures. To observe mitotic structures, we co-expressed MLKS2-mNeonGreen with pCaMV35S::AtCYCD3;1, which induces tobacco pavement cells to undergo mitosis (Xu et al., 2020). Like SUN1, MLKS2 can also localize to the preprophase band, phragmoplast, and new cell plate (Figures 7, B-D, Supplemental Movie S1).

### Polarized nuclear positioning, but not early polarity markers, are altered in *mlks2*

Proteins such as BRK1, PAN2 and PAN1 that are important for subsidiary cell polarization accumulate at the SMC-GMC interface, where the nucleus will eventually migrate (Figure 1A). Plants with mutations in genes encoding these proteins have defects in nuclear migration and polarized actin accumulation (Cartwright et al., 2009; Zhang et al., 2012; Facette et al., 2015). Since nuclear migration occurs after polar accumulation of BRK1, PAN2 and PAN1, we predicted there would be no effect on their polarized localization. However, it is plausible that polarized nuclei may feedback to these polarity markers, and influence their localization. As predicted, all of these polarity markers are correctly localized in *mlks2* (Supplemental Figure S8). Their polarized accumulation was maintained even after cell division in both wild type and *mlks2* (Supplemental Figure S8).

After the polarization of BRK1 and PAN proteins, the polarized accumulation of actin and nuclear migration to the future division site happen as concomitant steps (Facette et al., 2015). We predicted, based on the known functions of SUN-KASH proteins, that *mlks2* mutants might have a defect in nuclear migration to the future division site, prior to mitosis. To determine if nuclear position is altered in *mlks2* mutants, we used RanGAP-YFP, a nuclear envelope marker, to determine nuclear position in pre-mitotic SMCs. Nuclear polarization was categorized as: polarized (touching the cell wall and centered relative to the GMC); offset (touching the cell wall, but not centered relative to the GMC); or unpolarized (not touching the cell wall) (Figure 2A). We observed a significantly higher number of offset and unpolarized nuclei in *mlks2-1* (Figures 2, B and C).

**Figure 2.**
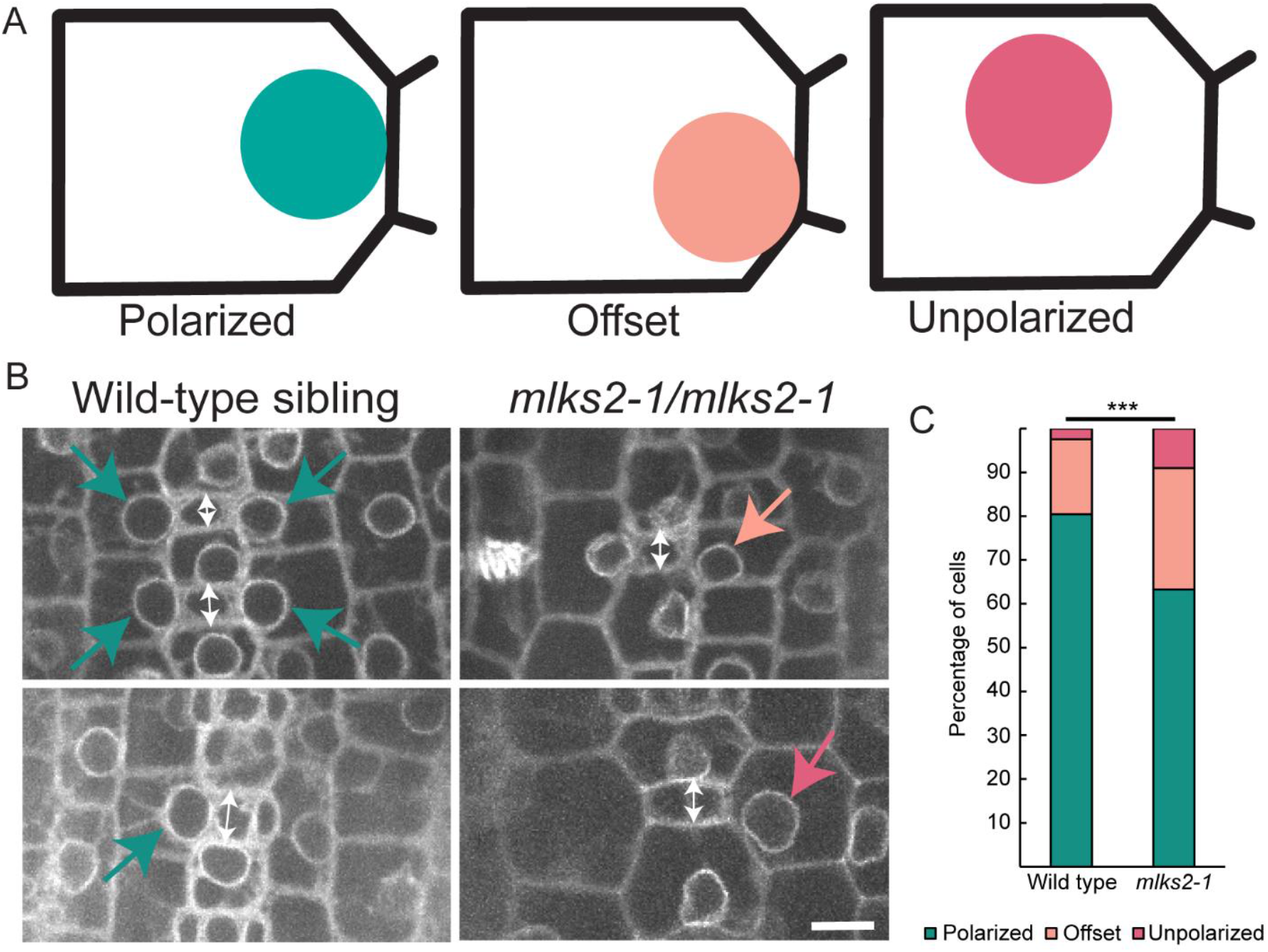
Nuclear position is impaired in *mlks2-1*. A, Nuclear polarization model (polarized, offset, and unpolarized) used for quantification. B, Visualization of nuclear position using RanGAP-YFP in wild type and *mlks2-1*. Double white arrowhead indicates the width of GMC and colored arrows, corresponding to (A), highlight the nuclear positioning. Scale bar = 10 μm. Same scale bar is applicable to all images. C, Quantification of the nucleus based on the polarization category in (A). Cells with GMC width 6μm or more were used for the quantification. 3 individual plants from both wild type and *mlks2-1* mutant were used for the experiment. 123 (polarized: 99, offset: 21, unpolarized: 3) and 155 (polarized: 98, offset: 43, unpolarized: 14) cells from wild type and *mlks2-1*, respectively, were observed for the quantification. The total number of cells based on nuclear polarization in each genotype are statistically tested by the Fisher’s Exact test. P value is 0.0004. ***P <0.001.

The role of the polarized actin patch is completely unknown. It is tempting to speculate that the actin patch, which sometimes has actin cables radiating towards the nucleus, plays a role in nuclear polarization. However, several different experiments using cytological and/or inhibitor studies suggest this is unlikely (Apostolakos et al., 2018). Mutations in *brk1, pan2* or *pan1* all affect both actin patch accumulation and nuclear migration, but on a cell-by-cell basis, actin patch formation and nuclear polarization can be uncoupled. To date, no mutant has been shown to have an effect on actin accumulation only, or nuclear polarization only. The *mlks2* mutant provides a good genetic tool to untangle the relationship (or lack thereof) between actin accumulation and nuclear position. If nuclear polarization and polarized actin accumulation are independent events, no effect on polarized actin accumulation is expected. However, if the nuclear position stimulates polarized actin accumulation, or the ARM domains of MLKS2 somehow stimulate actin patch formation independent of the nuclear position, actin accumulation at the polarized site might be abnormal. Using ABD2-YFP as a marker for actin, we quantified fluorescence intensity in *mlks2* mutants and wild type siblings at the polarized site and the adjacent cell wall in pre-mitotic SMCs, which is expressed as a ratio (Figures 3, A, B, and D). SMC nuclear positions were also classified as illustrated in Figure 3C. In both mutant and wild type SMCs, cells with polarized nuclei showed a higher fluorescence intensity (higher ratio) at the polarized site, but no differences in the ratio of intensities were observed in *mlks2* mutants, relative to siblings, regardless of nuclear position (Figure 3D). Importantly, we never observed a “delocalized” actin patch, which has been observed in *pan2* and *pan1* mutants (Zhang et al., 2012; Facette et al., 2015). Instead, in *mlks2* mutants, even when the nuclei were unpolarized or offset, the actin patch was always localized normally at the GMC-SMC contact site. These data demonstrate that MLKS2 functions in nuclear polarization but not actin patch formation, and that actin patch formation does not strictly depend on nuclear polarization.

**Figure 3.**
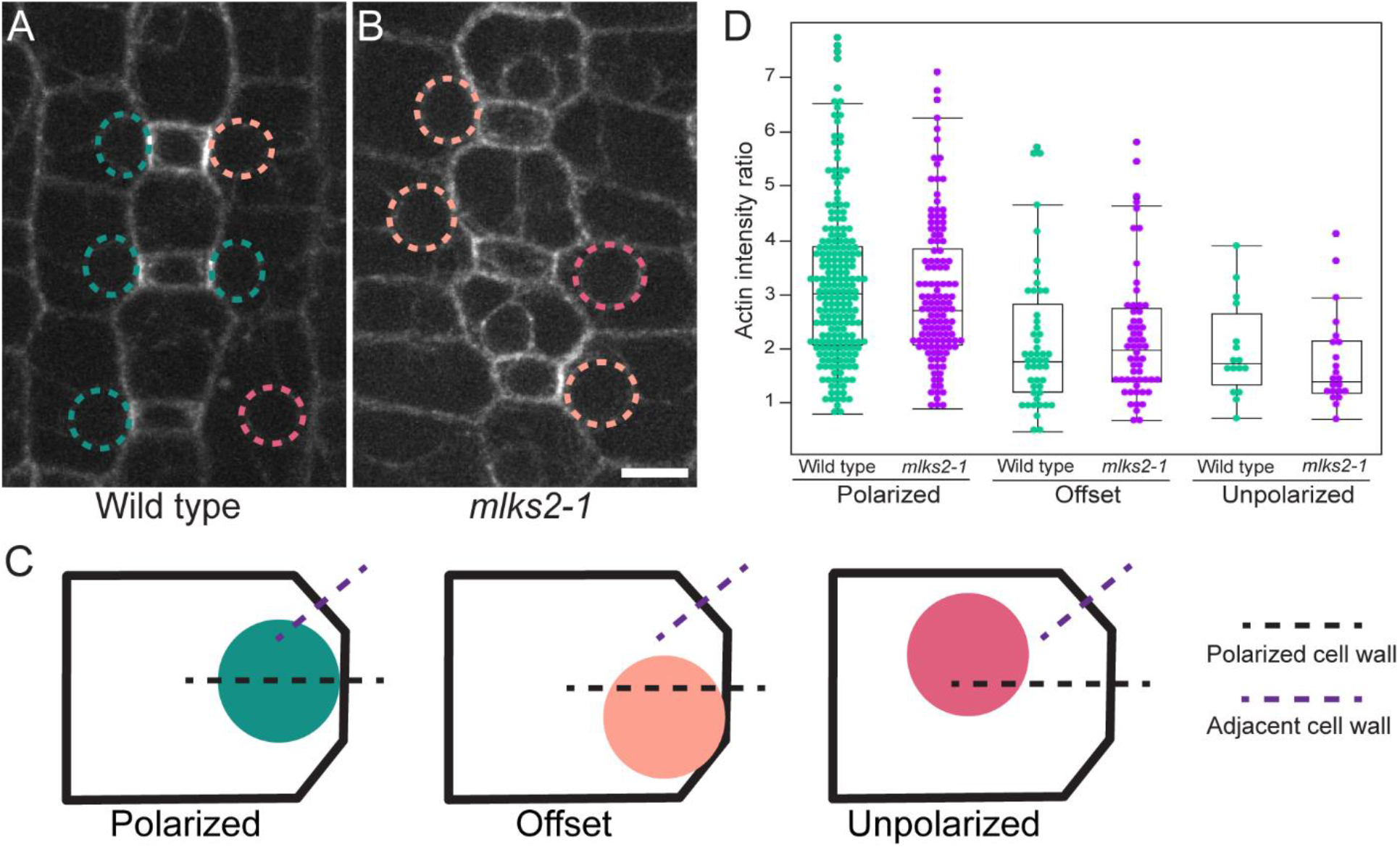
Actin patch formation remains unaltered in *mlks2-1*. Visualization of actin patch formation in the GMC-SMC interphase and nuclear position using ABD2-YFP in wild type (A) and *mlks2-1* (B). Nuclear position is marked with the colored dashed circle, corresponding to (C), for polarized, offset, and unpolarized nucleus. Scale bar = 10 μm. Same scale bar is applicable to all images. D, Actin patch quantification was based on comparing the intensities between polarized and adjacent cell walls. Actin intensity ratio (polarized/adjacent cell wall) in wild type and *mlks2-1* for three distinct categories of nuclear positioning. 3 individual plants of each genotype were used for the experiment. Cells with GMC width 5μm or more were used for the quantification. Total 270 (Polarized: 206, Offset: 48, Unpolarized: 16) and 206 (Polarized: 126, Offset: 58, Unpolarized: 22) cells from wild type and *mlks2-1*, respectively, were observed for the quantification.

### Preprophase bands are misplaced and spindles are affected in *mlks2*

In *mlks2* mutants, BRK1, PAN2, and PAN1 are polarized throughout mitosis and after division. Therefore, despite a defect in nuclear polarization, the cell can still perceive and maintain polarity in *mlks2*. What leads to abnormal division planes in *mlks2*? Abnormal nuclear polarization in *mlks2* may lead to abnormal division planes through the misplacement of mitotic structures. Alternatively, since MLKS2 can localize to mitotic structures after nuclear envelope breakdown (Supplemental Figure 7), MLKS2 may have a direct role in the assembly or function of mitotic structures. We used CFP-TUB to examine mitotic structures in *mlks2* and wild type siblings (Figures 4, A-C;5, A and B). No obvious structural deformation of the PPB (Figures 5, A-C) or spindle (Figures 5, A and B) was observed in *mlks2-1*. However, we observed a number of transverse PPBs in *mlks2-1* SMCs (Figures 4, A-C). 8.5% (38/446 cells) of SMCs had a misoriented PPB in *mlks2* vs. 2.1% (7/338 cells) in wild type siblings (Figure 4D). Notably, the percentage of SMCs with a misoriented PPB correlates with the observed percentage of abnormal subsidiary cells in both genotypes (Figure 1). A similar correlation between the proportion of misplaced PPBs and aberrant subsidiary cells has been described in the SMC polarity mutant *pan1* (Supplemental Figure S9) (Gallagher and Smith, 2000). To determine if there was a relationship between PPB and nuclear position, we again classified nuclear position, but only in cells that had a PPB (Figure 4D). Notably, many cells with correctly positioned PPBs had offset nuclei, in both wild type (7% of SMCs) and *mlks2* (9%). In *mlks2* there were more SMCs with offset nuclei, and about a third of these cells also had a misplaced PPB (Figure 4D). Cells with both an offset nucleus and misoriented PPB occurred just 0.9% of the time in wild type, but 5.6% of the time in *mlks2*. We also observed more cells in *mlks2* that had an unpolarized nucleus (0.8% in wild type, vs 2.0% in *mlks2)* and in both genotypes, these cells always had misplaced PPBs (Figure 4D). In summary, abnormal PPB placement in both *mlks2* and wild type SMCs is almost always associated with aberrant nuclear position.

**Figure 4.**
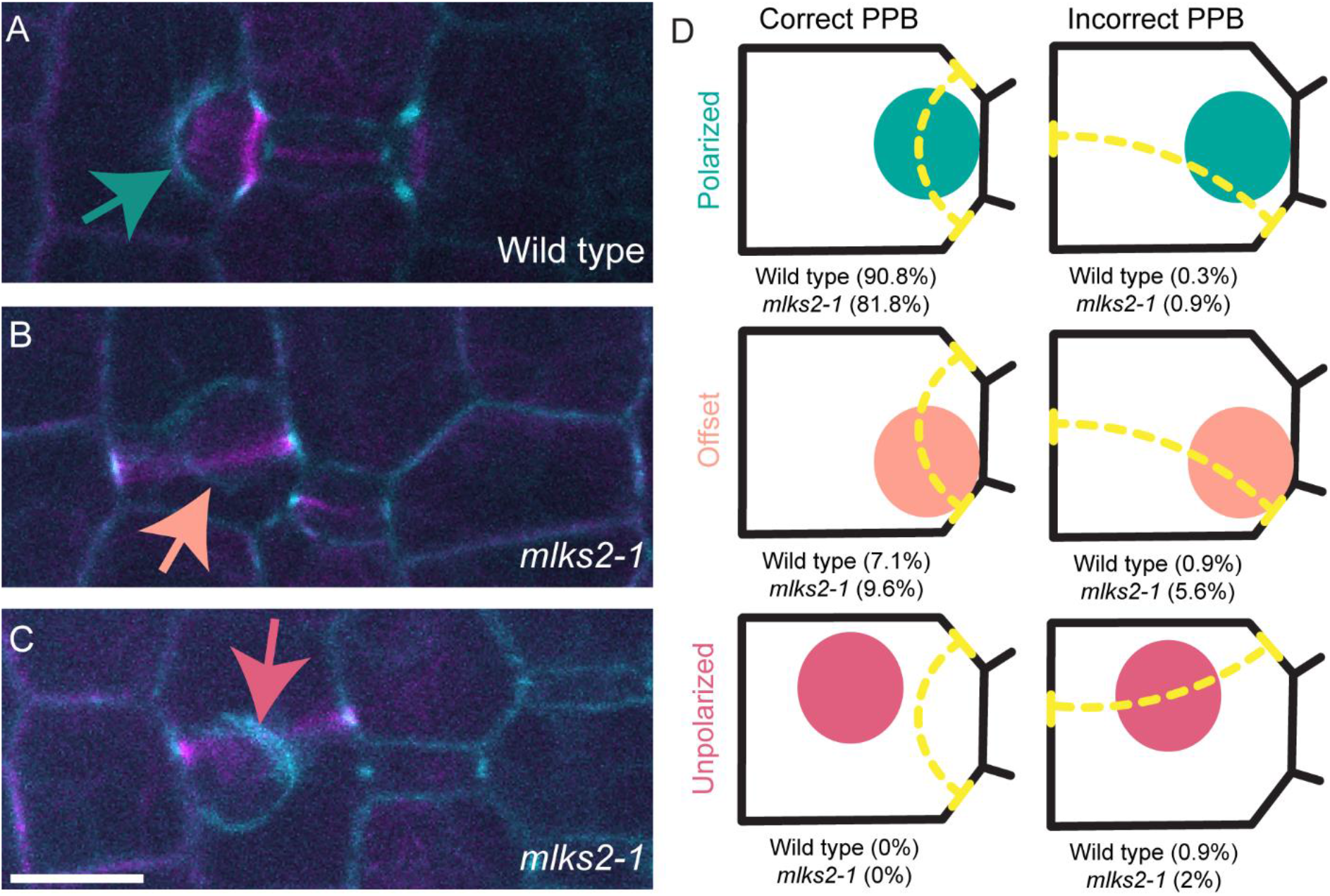
Nuclear positioning defect leads to misplaced PPB. Observation of PPB formation and nuclear position from cortex (magenta) and midplane (cyan), respectively, using CFP-TUB in wild type (A) and *mlks2-1* (B, C). Colored arrow indicates the nuclear position corresponding to (D). Scale bar = 10 μm. Same scale bar is applicable to all images. D, Six probable cellular events combining PPB placement (correct/incorrect) with nuclear position (Polarized/Offset/Unpolarized). Percentage of cells observed for corresponding events are mentioned in for both wild type and *mlks2-1*. 3 individual plants of each genotype were used for the experiment. Total 338 and 446 cells from wild type and *mlks2-1*, respectively, were observed for the quantification.

**Figure 5.**
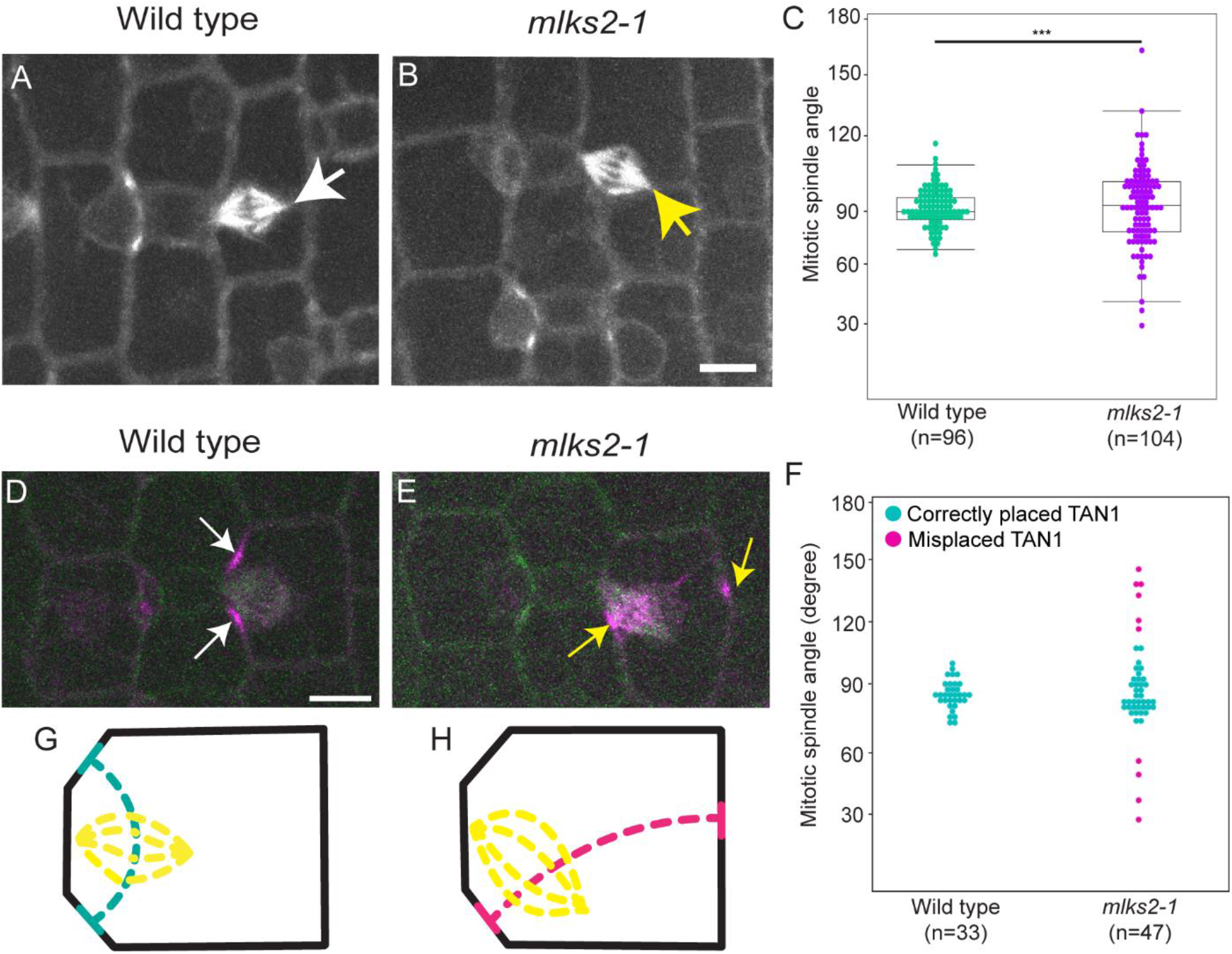
Mitotic spindle angle depends on the pre-determined division site in the *mlks2-1*. Mitotic spindle morphology is observed using CFP-TUB for wild type (A) and *mlks2-1* (B). White arrow indicates the mitotic spindle with an angle close to 90°. Yellow arrow indicates the mitotic spindle deviated from ideal 90°. Scale bar = 10 μm. Same scale bar is applicable to all images. C, Spindle angle was quantified. At least 3 plants from each genotype are used for the experiment. 96 wild type and 104 *mlks2-1* cells were considered for the quantification. The total number of cells judged statistically based on mean (P=0.5169) and variance (P<0.0001) by 2-sided F-test. ***P <0.0001. Representative images of mitotic spindle and cortical division site using CFP-TUB (green) and TAN1-YFP (magenta), respectively, in wild type (D) and *mlks2-1* (E). Scale bar = 10 μm. Same scale bar is applicable to all images. F, Quantification of mitotic spindle angle in wild type and *mlks2-1*. Mitotic spindle angle from the cells with correctly placed and misplaced TAN1 are marked with cyan and magenta dots, respectively. Graphical model for division site-based spindle angle orientation in wild type (G) and *mlks2-1* (H).

We also measured the mitotic spindle angle. Only spindles where one end was anchored at the GMC-SMC interface were measured. Although the mitotic spindle angle has a central tendency towards 90^0^ in both wild type and *mlks2-1*, the mitotic spindle angle varies within a significantly larger range (30^0^-170^0^) in *mlks2-1* (Figures 5, A-C). This variation in angle could be caused by multiple mechanisms, which are not mutually exclusive. The spindle typically orients perpendicularly to the axis of the specified cortical division site, as marked by the PPB and other division site markers. The increased frequency of misaligned spindles could be a result in an increased variation in the angle of the cortical division site. Alternatively, the spindle itself could be “wobbly”, and rotate during mitosis. To test this hypothesis, we further investigated spindle positioning in dividing SMCs, but this time we measured all spindles – whether they were anchored at the GMC or in the center of the cell. We measured spindle angle in cells by co-expressing the cortical division site maker TAN1-YFP with CFP-TUB (Figures 5, D-F). In *mlks2* mutant, the more extreme spindle angles were always observed in cells with TAN1-YFP that marked an oblique division site atypical of polarized SMCs (Figures 5, D and E). This data indicates that mitotic spindle orients based on the PPB placement and future division site marking (Figures 5, G and H).

### The mitotic apparatus follows the initial division site marked by PPB

To determine the precise relationship between nuclear position, preprophase band placement, and final division plane, we performed live-imaging of SMCs undergoing mitosis. We had several important questions to address using our live imaging. Firstly, how is the position of the subsequent mitotic structures, the spindle and the phragmoplast, influenced by nuclear position and/or the PPB in *mlks2*? Secondly, what is the relationship between PPB position and nuclear position? Can the PPB influence nuclear position? Thirdly, are offset and unpolarized nuclei mispositioned because their initial guidance towards the guard mother cell fails, or because their nuclear position at the division site is unstable? That is, how exactly does MLKS2 influence nuclear position?

We addressed these questions using both HIS-YFP and CFP-TUB markers in *mlks2-1* and wild type siblings. Histone marks the nuclear position and helps to identify the transition from G1 to G2 phase, while CFP-TUB facilitates the visualization of cell division structures and events, such as the PPB, spindle, phragmoplast, and cytokinesis. We imaged the SMC divisions occurring in the stomatal division zone of leaf 4 in wildtype siblings and *mlks2* plants at 2-minute intervals (Figure 6A and Supplemental Movies S2 and S3). We observed nuclear position and complete cell divisions (PPB to cytokinesis) in 62 cells (27 wild-type and 35 *mlks2-1* cells). In wild type cells, all of the PPBs were correctly positioned, and in 26/27 cells the nucleus was polarized when the PPB was present (Figure 6B). Mid-plane views of mitosis and cytokinesis show that after PPB disassembly, the spindle forms perpendicular to the PPB, followed by the phragmoplast, which forms at the expected division site (Supplemental Movie S2). In the *mlks2* mutant, 10/25 cells had a misplaced PPB, and in all 10 of these cells the nucleus was either offset (6 cells) or unpolarized (4 cells) (Figure 6B). In all cases, the spindle and phragmoplast orientation matched the orientation of the PPB (Figure 6B). For example, Supplemental Movie S3 shows a *mlks2* SMC where the nucleus is initially offset, and the PPB is transverse to the axis of the cell. The spindle shows an oblique angle, and the phragmoplast re-aligns with the former PPB position.

**Figure 6.**
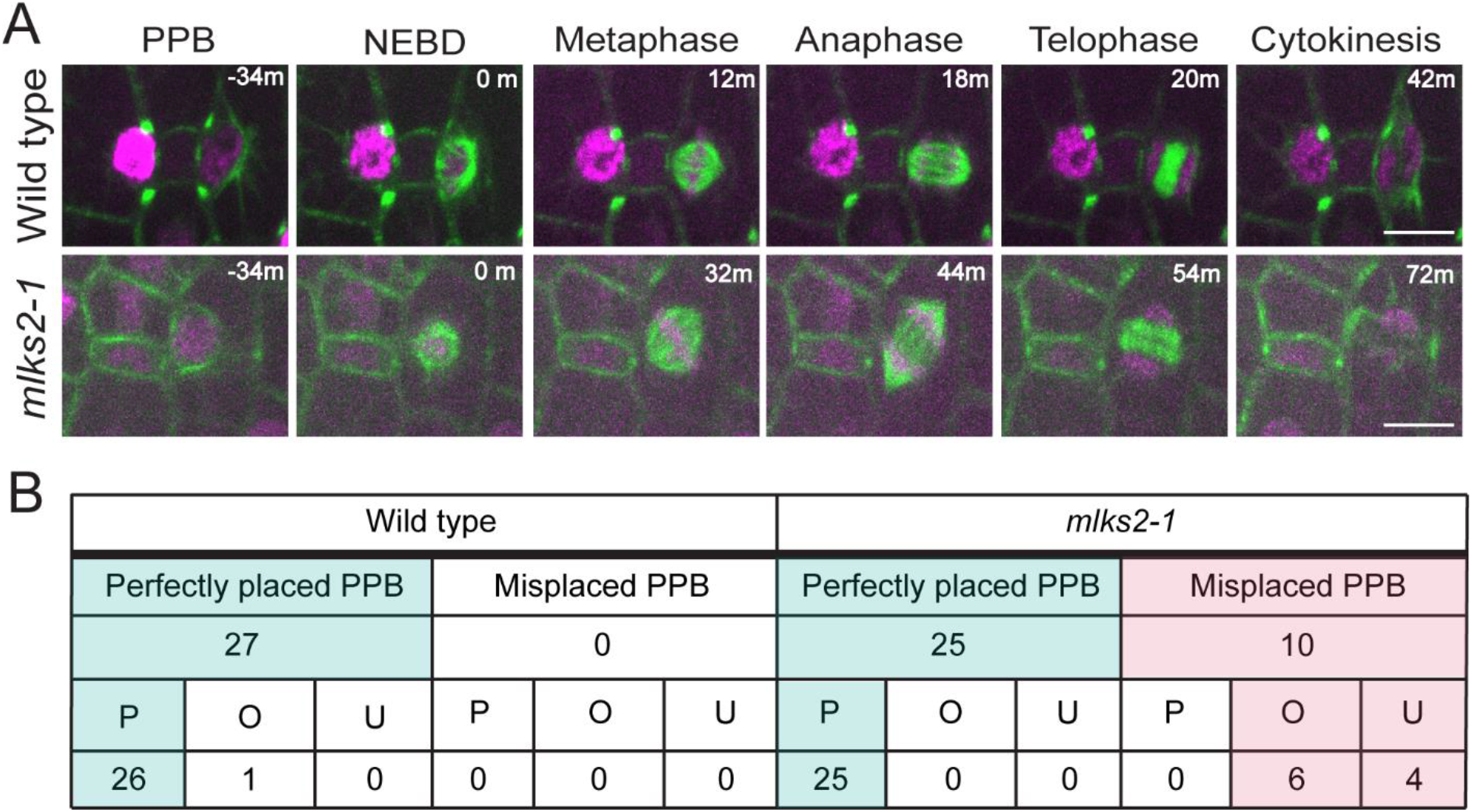
Mitotic apparatus follows the division site marked by PPB. A, Timelapse of asymmetric cell division during subsidiary cell formation in wild type (upper panel) and *mlks2-1* (bottom panel). The nucleus and microtubules were visualized using HIS-YFP (magenta) and CFP-TUB (green), respectively. Nuclear envelope breakdown is considered as the starting point and marked as 0 min. Timelapse movies were captured in every 2 min interval. Representative images highlighting key cell division stages: PPB, NEBD, metaphase, anaphase, telophase, and cytokinesis were presented. Scale bar = 10 μm. Same scale bar is applicable to all images. B, Quantification of PPB placement (Perfectly placed PPB/Misplaced PPB), nuclear positioning (P-polarized; O-offset; U-unpolarized), and cell division orientation (Expected division plane/Misoriented division plane) from the timelapse observation. Perfectly placed PPB and misplaced PPB categories are highlighted with distinct color in the table.

**Figure 7.**
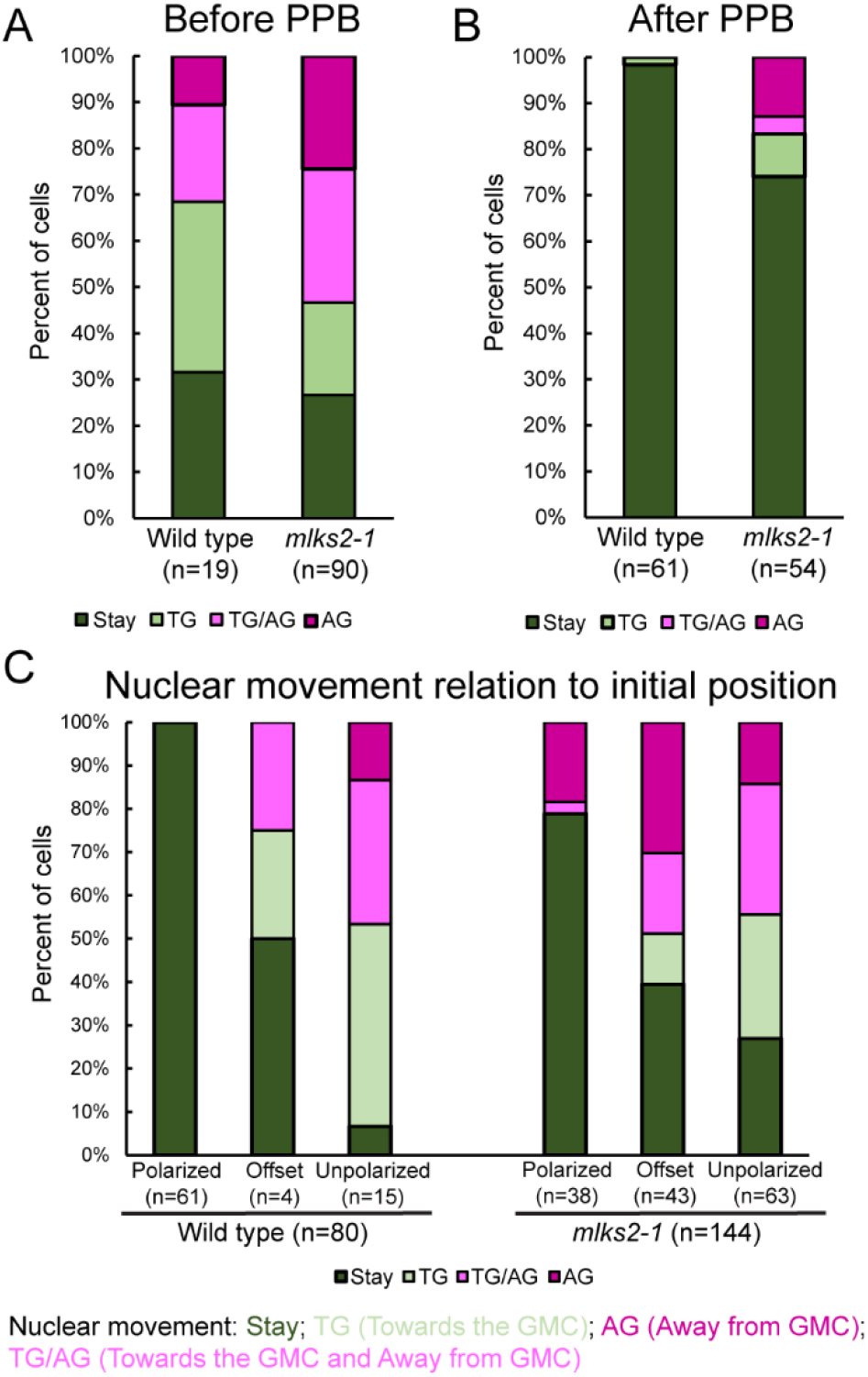
Nuclear migration and anchorage are defective in *mlks2-1*. Analysis of nuclear movement in wild type and *mlks2-1* in the absence (A) and presence (B) of the PPB. Nuclear movements were divided into 4 categories: “stay”, “towards the GMC (TG)”, “away from GMC (AG)”, and oscillating movement “towards the GMC and away from GMC (TG/AG)”. (C) The same data as in A and B, but instead classified by initial nuclear position rather than PPB presence/absence. In all cases, the nucleus is considered to have moved in a particular direction if it is displaced more than one third of the nuclear width. A total of 80 cells from 8 wildtype plants and 144 cells from 5 *mlks2* plants were analyzed.

We measured the total time for mitosis in the entire population of *mlks2* SMCs, and found that metaphase and anaphase timing are similar between perfectly oriented and misoriented cell divisions (Supplemental Figure S10A), but telophase and cytokinesis sometimes require a longer time during misoriented cell divisions (Supplemental Figure S10B). This longer time is presumably due to the longer path covered by a transversely formed phragmoplast (Supplemental Figures S10, D and E). When the time taken for telophase and cytokinesis is normalized by the distance covered by phragmoplast, there is no difference between wild type and *mlks2* (Supplemental Figure S10C). Together, these data suggest that spindle and phragmoplast position are not directly impaired in *mlks2* SMCs, but rather their misalignment is a result of aberrant PPB positioning.

### MLKS2 is required for both nuclear migration and anchorage

What is the relationship between PPB position and nuclear position? As suggested by our static image data and Supplemental Movie S3, nuclear position influences PPB position. But can the PPB influence nuclear position, and how stable is nuclear position after PPB formation? To answer these questions, we used timelapse imaging to observe nuclear movement in the stomatal division zone (Supplemental Movie S4). All SMCs that did not divide were categorized by the presence/absence of the PPB, the initial nuclear position, and the directional movement of the nucleus. Movement was categorized as towards the GMC (TG) (Supplemental Movie S5), away from GMC (AG) (Supplemental Movie S6), moving both towards and away from the GMC (TG/AG) (Supplemental Movie S7) or unmoving (stay) (Supplemental Movie S8). The nucleus was considered to have moved if it was displaced more than half of the nuclear width. The data is summarized in Figure 7 and Supplemental Movies S5-S10. In both mutant and wild type cells, more movement is observed prior to PPB formation, with the nucleus becoming quite stable after PPB formation (Supplemental Movie7) (Figure 7A,B). This analysis is difficult and time-intensive, but we observed certain behaviors in *mlks2* that we never or seldom observed in wild type. For example, after PPB formation, the nucleus never moved away from the polarized site in wild type cells (Supplemental Movie S8). However, we frequently saw unpolarized nuclei, or nuclei that moved away from the GMC after PPB formation in *mlks2* (Supplemental movies S9, S10). In fact, regardless of PPB presence, once a wild type nucleus touched the wall of the GMC, it never moved away – but we also observed the nucleus move away from the polarized site in *mlks2* cells prior to PPB formation, which we never observed in a wild type cell (Supplemental Movie S6). Our data indicate that initial migration might be affected, and maintenance of nuclear position is definitely affected in *mlks2* SMCs.

## Discussion

Together, our findings shed new light on the function of MLSK2 in asymmetric cell division and generated new insights into the role of nuclear position in division plane determination. Our data indicates that nuclear position during volumetrically asymmetric divisions is important for establishment of the cortical division site (Figure 8). Positional cues from the nucleus feed into the orientation of the PPB, which then guides the orientation of the subsequent mitotic structures. Aberrant nuclear positioning results in misplaced PPBs and division planes (Figure 8).

**Figure 8.**
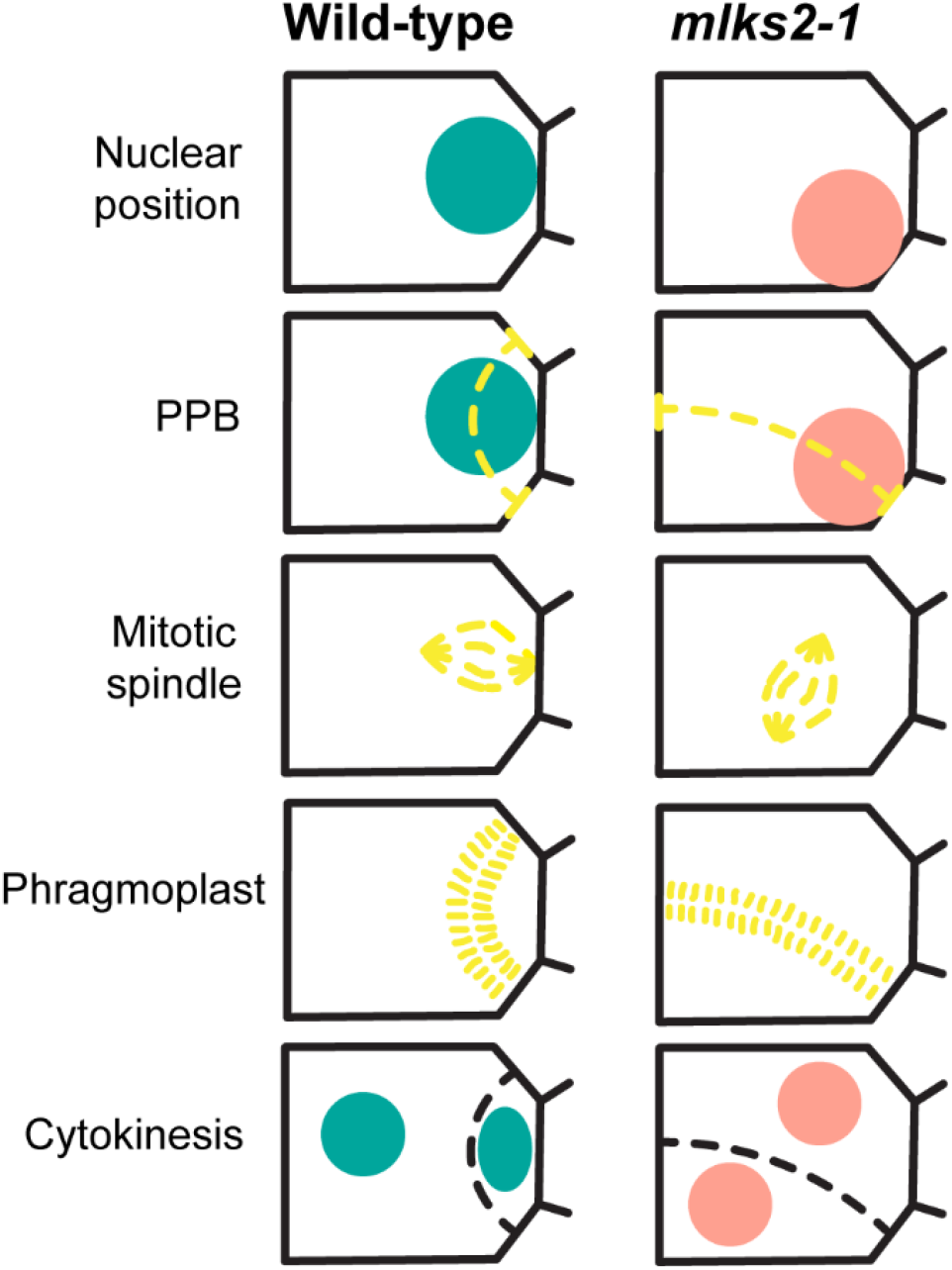
Graphical model of outer nuclear membrane protein-mediated nuclear positioning and future division site determination. (Left) In wild type condition, nucleus polarizes based on the established cell polarity, perfect PPB placement, and the mitotic apparatus follows the division site marked by PPB until cytokinesis. (Right) In *mlks2-1*, nucleus fails to position according to the established cell polarity, misplaced or transverse PPB is formed based on the impaired nuclear position, and the mitotic apparatus follows the division site marked by PPB until cytokinesis.

Our data illustrates several key findings. Firstly, the nuclear envelope protein MLKS2 is important for a polarized nuclear position before and after formation of the PPB. Once the PPB forms, division proceeds normally. However, in a subset of cells, the nucleus was displaced at the onset of mitosis, even in otherwise polarized cells, which results in an abnormal specification of the division site and subsequent division. Notably, many cells still had correctly polarized nuclei. This could be for several reasons. MLKS2 has an orthologue, MLKS1 (Gumber et al., 2019b). Double mutants may produce a more dramatic phenotype. Other nuclear envelope complexes, such as the nuclear pore, may also contribute to cytoskeletal interactions and nuclear positioning (Best et al., 2021). Previous inhibitor studies in maize indicate that pre-mitotic nuclear migration is driven by actin networks, while maintenance of a polarized position is driven by microtubule networks (Panteris et al., 2006). MLKS2 has been shown to co-localize with actin (Gumber et al., 2019a), but it is plausible it may also interact with the microtubule cytoskeleton, directly or indirectly. Future studies determining the physical and phenotypic connections between LINC complex proteins and specific cytoskeletal networks will help determine the mechanism of nuclear migration and maintenance of position. In animal cells, LINC complex proteins have been found to interact with the actin cytoskeleton and microtubule motors (Burke, 2019).

A second key finding is that perturbation of nuclear position results in a misplaced PPB and subsequent asymmetric divisions, even in the presence of polarization cues. The misplaced PPBs typically took on a consistent, transverse orientation, which is perpendicular to the long axis of the leaf. Symmetric divisions of maize leaf epidermal cells typically display transverse divisions (Martinez et al., 2018), suggesting the SMCs may be responding to a broader cue such as cellular shape, or an unknown cue. Interestingly, many of the aberrant divisions took place in mutant cells where the nucleus was only slightly offset from the polarized position. In both wild type and mutant cells, offset nuclei occur 7-9% of the time, with completely normal PPB positioning. Why, in the mutant, do offset cells sometimes result in aberrant division site specification, while in other cases it is normal? High resolution imaging of the nuclear envelope and cytoskeleton at the time of preprophase band establishment may help resolve this question. Indeed, understanding the events at the nuclear interface in general, may help resolve how exactly the nucleus, perhaps through nucleating events, promotes PPB positioning.

Other studies have called into question the role of nuclear position and even PPB formation, in division plane determination. How can these studies be reconciled with our finding that nuclear polarization is key for proper orientation of asymmetric divisions via its effect on PPB placement? Previous studies demonstrated that nucleus is connected to PPB through microtubule radiating from the nuclear envelope and in the presence of offset PPB, nucleus moves to align with the offset PPB (Granger and Cyr, 2001). Notably, in Arabidopsis roots that lack a PPB through genetic manipulation, division planes are relatively normal (POK1 localization), as is overall tissue patterning (Schaefer et al., 2017). Formation of different cell layers in the Arabidopsis root occurs via asymmetric division, however these developmentally asymmetric divisions create volumetrically similarly sized cells (Facette et al., 2019). In these asymmetric divisions, the nucleus fills much of the cell, and nuclear polarization is not a prominent feature. Therefore, cues independent of the nucleus and the PPB, including cortical polarity and mechanical cues, may be sufficient to ensure developmental asymmetry and proper orientation of division. Another case where our findings may seem to disagree with previous observations is that caulonema tip cells of *Physcomitrella patens* execute oriented divisions without forming PPBs. However in this case, cortical proteins such as PpKin12-Ie and PpREN (homologs of AtPOKs and PHGAP1/2, respectively), mark the division site prior to completion of cytokinesis (Miki et al., 2014; Yi and Goshima, 2020). We speculate that pre-mitotic nuclear position may play a key role in localizing cortical division site markers in cells lacking PPBs. As exemplified by these two cases, division plane establishment is an essential and complex process that likely has many combinatorial factors, therefore the context of cell division must always be considered. In volumetrically asymmetric divisions, such as in the SMC, nuclear position and PPB establishment appear to be especially important.

## Materials and Methods

### Plant materials

Stably expressed fluorescent marker lines: BRICK1-CFP (Facette et al., 2015), PAN2-YFP (Zhang et al., 2012), PAN1-YFP (Humphries et al., 2011), FIMBRIN (ABD2-2xYFP) (Mohanty et al., 2009), CFP-TUBULIN-BETA (Mohanty et al., 2009), TUBULIN-YFP (Mohanty et al., 2009), HISTONE H1B-YFP (Mohanty et al., 2009), TANGLED1-YFP (Mohanty et al., 2009), RanGAP-YFP (obtained from MaizeGDB stock center; UWYO-FP032) were described previously (Krishnakumar et al., 2015). *mlks2-1* (mu1038603/UF-Mu04133) and *mlks2-2* (mu1058535/UF-Mu07312) mutant seeds were a gift from Hank Bass (Florida State University). *pan1-ems* was provided by Laurie Smith (University of California San Diego) and described previously (Gallagher and Smith 2000). p35S::AtCYCD3;1 plasmid was generously shared by Bo Liu (University of California Davis) (Xu et al., 2020). Single and double fluorescent protein marker lines with *mlks2-1* and *mlks2-2* were generated through genetic crosses.

### Plant growth condition

Plants used for experiments were grown for approximately 2 weeks in the greenhouse condition (temperature between 20°C to 29°C and 16h day length). Plants were grown in Pro-mix Professional soil supplemented with Peters Excel 15-5-15 CalMag fertilizer and chelated liquid iron (Southern Ag). Supplemental LED lights (Fluence VYPR series) were used to maintain a 16-hour day length. For the field sample, plants were grown in the farm facility of the University of Massachusetts Amherst in the summer of 2021. For all experiments, plants were grown in parallel and wild type siblings were used as controls.

### Stomatal phenotyping

Fully expanded leaf 3, 5, 7, and 10 were used for glue impression (Allsman, Dieffenbacher, and Rasmussen 2019) and prepared for the phenotyping slide. These slides were observed for stomatal phenotype using Nikon SMZ800N attached with ED plan 2X WF WD:35 objective lens and coupled with AmScope camera MU1003.

### Stomatal conductance measurement

Stomatal conductance measurements were performed using CIRAS3 Portable Photo-Synthesis System. The leaf cuvette will maintain a stable environment: CO_2_ concentration is 410 μmol mol^−1^; H_2_0 reference is 80%; the cuvette flow is 300 cc min^-1^. For all the measurements, healthy and fully expanded leaf 3 were used. Briefly, the leaf cuvette clamped the leaf and waited at least 30 mins to stabilize the initial stomatal conductance under the light (1000 PPFD). Lights were turned off immediately after the initial conductance became stable, and the system recorded the transpirational conductance of the leaf surface simultaneously for the whole period of treatment. The data presented are the mean ± SE of at least 4 individual plants. Relative gs was computed for each individual measured plant by normalizing gs to the steady initial gs value observed. Data analysis was performed in R and JMP^®^ 16.1.

### Molecular cloning

Coding sequences of MLKS2 (primer pair: AA106 and AA107) and mNeonGreen (primer pair: AA108 and AA109) were amplified from *Zea mays* B73 cDNA and mNeonGreen-2A-mTurquoise2 (Addgene #98885), respectively. These amplified fragments are Goldengate compatible for type IIs restriction enzyme *Aar I*. Both MLKS2 and mNeonGreen were assembled into pType IIs plasmid using GeneArt™ Type IIs Assembly Kit, *Aar I* (Thermo Fisher Scientific #A15916). MLKS2-mNeonGreen from the pType IIs vector was further amplified (primer pair: AA98 and AA47) using attB1 and attB2 sites containing primers. This amplified product was first cloned into the entry vector pDONR221 using Gateway™ BP Clonase™ II Enzyme mix (Thermo Fisher Scientific #11789020) and eventually into the binary vector pMDC32 (ABRC #CD3-738) using Gateway™ LR Clonase™ II Enzyme mix (Thermo Fisher Scientific #11791020). Each cloned plasmid was confirmed by both restriction enzyme digestion and Sanger sequencing.

### Transient expression and mitotic induction in *Nicotiana benthamiana*

For transient expression in *Nicotiana benthamiana, Agrobacterium* strain GV3101 harboring different constructs were resuspended in infiltration buffer (10 mM MES pH 5.7), 10 mM MgCl_2_, 50 mg/L acetosyringone), and adjusted OD600 to 1.0. Equal volumes of cultures containing different constructs were mixed for co-infiltration. The resulting cultures were infiltrated into leaves of 3-to 4-week-old *Nicotiana benthamiana* plants. Leaf samples were imaged 48 hours after infiltration.

For mitotic induction, equal volumes of cultures containing different constructs (pCaMV35S::ZmMLKS2-mNeonGreen and pCaMV35S::AtCYCD3;1) were mixed for co-infiltration. Leaf samples were imaged 48 hours after infiltration.

### Confocal microscopy

Plant tissues were observed using a custom-built spinning disc confocal unit (3i) equipped with an inverted fluorescence microscope (IX83-ZDC, Olympus) CSU-W1 spinning disc with 50 micron pinholes (Yokogawa), a Mesa Illumination Enhancement 7 Field Flattening unit (3i), an Andor iXon Life 888 EMCCD camera and a UPLANSAPO ×60 Silicone Oil-immersion objective (NA = 1.20, Olympus) and 4 laser stack with TTL controller (3i). For CFP, YFP/mNeonGreen and RFP imaging of maize and tobacco plants, a 445/515/561 dichroic (Chroma) was used. All emission filters are from Semrock. A 445 nm laser line and 483/45 emission filter (CFP) or 514 nm laser and 542/27 emission filter (YFP/mNeonGreen) was used.

### Time-lapse imaging

For the time-lapse imaging, the stomatal division zones of leaf 4 of corresponding plants were dissected and mounted with water. Cell division events were captured every 2 minutes with the spinning disk confocal microscope at Facette lab (details mentioned in the confocal microscopy section) for 50-180 minutes.

### Image processing and quantification

Nuclear position (Figure 2), actin patch formation (Figure 3), correlation between PPB placement and nuclear position (Figure 4), spindle angle measurement (Figure 5), and nuclear movement categorization (Figure 7) were quantified in this study. The quantification strategies are explained in the following sections.

### Nuclear positioning quantification in static images

RanGAP-YFP marks the nuclear envelope and was used as a tool to observe the position of the nucleus. Previous study suggested that actin patch/nuclear polarization happens at the developmental stage, when the GMC width is 6μm (Facette et al. 2015). The stomatal division zone of L3 of both wild type and *mlks2-1* plants were used to image RanGAP-YFP and SMCs associated with GMC width 6μm or more were considered for the nuclear position quantification. As shown in Figure 2A, If the nucleus is perfectly localized with the polarized cell wall, it is considered as a “polarized” nucleus. If the nucleus is not perfectly polarized, but touches an adjacent cell wall of the polarized cell wall, then it is considered as an “offset” nucleus. Finally, if the nucleus is far away from both the polarized and adjacent cell wall, then it is considered as an “unpolarized” nucleus. Based on this explained and demonstrated nuclear positioning rule, we have categorized nucleus into: Polarized, Offset, and Unpolarized.

### Actin patch quantification

ABD2-YFP marker line was utilized to observe actin and nucleus position. Previously, it has been demonstrated that actin patch forms at the developmental stage, when the GMC width is 6μm (Facette et al. 2015). For our quantification, we have considered SMCs adjacent with GMC width 5μm or more. First, the actin patch was quantified from both polarized cell wall and adjacent cell wall by drawing a straight line at ImageJ/Fiji (Figure 3C). Then the intensity values were converted into ratios (polarized/adjacent) for each cell. These calculated intensity ratios were plotted at the Y-axis of Figure 3D. During the actin patch quantification, nucleus positioning was considered from the same cell. For the nucleus position quantification, similar rules were applied as RanGAP-YFP (Figure 2A).

### Correlation between PPB placement and nuclear position

To observe both nuclear position and PPB, we have used the CFP-TUB marker line. CFP-TUB clearly marks PPB and nucleus from midplane and cortical PPB ring from the cortex. Cells with PPB were only considered for this quantification. If a SMC contains the PPB, then nuclear position in the same cell is observed, based on the rule explained in the earlier sections (Figure 2A).

Image processing was performed using ImageJ/Fiji and Adobe Illustrator version 26.0 for only linear adjustments and preserving hard edges.

### Spindle angle measurement

To visualize the spindle angle, CFP-TUB was used as marker for the visualization. Cells with mitotic spindles were considered for the analysis. Mitotic spindle is supposed to create a right angle with the polarized or GMC cell wall. To measure the angle, one line was drawn at the polarized or GMC cell wall and another line was drawn from the pointed end of mitotic spindle towards the polarized or GMC cell wall. For figure 5A-C, the mitotic spindle anchored one end correctly was measured.

Spindle angle was measured in presence of cortical division marker TAN1-YFP (Figure 5D-F). Mitotic spindles in presence of TAN1-YFP have been considered for the analysis. Similar spindle angle measurement approach has been implied like Figure 5A-C, and spindles with either one end attached to the polarized cell wall or both end away from the polarized cell wall was considered for the analysis.

### Quantification of nuclear position in time lapse images

Nuclear movement has been observed using HIS-YFP along with CFP-TUB. Nuclear movement has been observed before and after the PPB formation. If the nuclear displacement is more than half of the nuclear width, then it has considered as nuclear movement. Nuclear movement from the timelapse movie has been divided into four categories: stay, towards the GMC (TG), away from GMC (AG), and towards the GMC and away from GMC (TG/AG). Usually after the nuclear polarization and PPB formation, the nucleus has a stable position. In this context, the nucleus was marked as “stay”. Before PPB formation, nucleus polarizes to the GMC and this nuclear movement is categorized as “towards the GMC (TG)”. If nucleus moves far away from the GMC, it has been categorized as “away from GMC (AG)”. If the nucleus moves in both directions (towards the GMC and away from GMC), this nuclear movement has been marked as “towards the GMC and away from GMC (TG/AG)”. This rule has been followed to categorize nuclear movement from timelapse movies and generate graph in Figure 7A-C. In the case of Figure 7C, the initial nuclear position has been used as a sub-category. For nuclear positioning, a similar rule as illustrated in Figure 2A has been applied.

### Statistical analysis

Statistical tests, p value, number of plant samples, and number of individual cells used for experiments and quantification are mentioned in the corresponding figure legends. Data acquisition was done in Microsoft Excel; plotting the data and statistical tests were performed using JMP^®^ 16.1.

### Accession numbers

Gene and protein sequences for maize and Arabidopsis can be found at MaizeGDB (www.maizegdb.org) and TAIR (https://www.arabidopsis.org/index.jsp), respectively. Accession numbers for maize Version 4.0/Version 5.0 of the B73 genome are:

*Mlks2*: Zm00001d052955 / Zm00001eb034900; *Pan1*: Zm00001d031437 / Zm00001eb200470; *AtCYCD3*: At4g34160.

## Acknowledgments

We would like to thank Hardeep Gumber (Florida State University) and Hank Bass (Florida State University) for their initial work on the maize LINC complex and providing seeds. Authors are grateful to Bo Liu (University of California Davis) for providing p35S::AtCYCD3;1 plasmid for the mitotic induction experiment. Erika Norris (Undergraduate student at Facette lab) counted aberrant cells from summer field samples. The color palette for drawing model and illustration in figures has been inspired and taken from Wes Anderson’s films.

## List of Supplemental data

### Supplemental Figures

**Supplemental Figure S1**. MLKS2 is an outer nuclear membrane protein and part of the LINC complex.

**Supplemental Figure S2**. MLKS2 is required for proper interstomatal cell and guard cell division.

**Supplemental Figure S3**. MLKS2 regulates the first asymmetric cell division.

**Supplemental Figure S4**. Different abnormal divisions result in different stomatal phenotypes in *mlks2*.

**Supplemental Figure S5**. Aberrant subsidiary cell frequency of *mlks2-1* varies based on growth condition.

**Supplemental Figure S6**. Stomatal conductance is not affected in *mlks2-1*.

**Supplemental figure S7**. MLKS2 localizes at the nuclear membrane and with mitotic apparatus.

**Supplemental Figure S8**. Earlier cell polarity markers remain the same in *mlks2-1*.

**Supplemental Figure S9**. Transverse PPB formation in the *pan1-ems*.

**Supplemental Figure S10**. Mitosis timing is not altered in *mlks2-1*.

### Supplemental Movies

**Supplemental Movie S1**. MLKS2 localizes at the nuclear membrane and with phragmoplast and new cell plate. Transient tobacco expression with pCaMV35S::MLKS2-mNG and pCaMV35S::AtCYCD3. Time lapse movie was captured after 48h of the transformation. The images were captured in every 2-minute interval.

**Supplemental Movies S2 – S10:** All Movies are played at 5 frames per second, with frames taken at 2-minute intervals. In all movies CFP-TUB is green and HIS-YFP is magenta.

**Supplemental Movie S2**. Normal asymmetric division of an SMC in a wild type sibling.

**Supplemental Movie S3**. Abnormal transverse division of an SMC in *mlks2-1*.

**Supplemental Movie S4**. Overview of the stomatal division zone in *mlks2-1*. The video shows 3 stomatal rows and a variety of nuclear behaviors.

**Supplemental Movie S5**. “TG” nuclear movement in a wild type SMC before PPB formation. Normal asymmetric division of an SMC in a wild type sibling is shown in the top most SMC. In the bottom SMC, PPB formation has not yet occurred and the cell has an unpolarized nucleus with a stable position. The nucleus moves towards the GMC at the end of the video. Classified as “TG”.

**Supplemental Movie S6**. “AG” nuclear movement in a *mlks2-1* SMC before PPB formation. The nucleus is initially polarized and touches the SMC wall closest to the GMC. The nucleus moves away and touches the side wall and eventually the wall distal to the GMC.

**Supplemental Movie S7**. “TG/AG” nuclear movement in a *mlks2-1* SMC before PPB formation. The nucleus is initially unpolarized, and then becomes offset, then polarized, and then moves away from the GMC to become unpolarized.

**Supplemental Movie S8**. Stable polarized nuclear positioning in wild type SMCs after PPB formation. Six SMCs are shown. The SMC in the bottom left is dividing, all other SMCs have PPBs and the nuclear position is stable. These nuclei are classified as “stay”.

**Supplemental Movie S9**. Unpolarized nuclei in SMCS after PPB formation. Six SMCs are shown. The cell in the bottom right divides normally. All other SMCs have a PPB. Two nuclei are initially offset and move towards and away from the GMC.

**Supplemental Movie S10**. “TG/AG” nuclear movement in a *mlks2-1* SMC after PPB formation. An unpolarized nucleus is initially unpolarized and shows movements both towards and away from the GMC.

### Supplemental Tables

**Supplemental Table S1**. Primers used in this study for genotyping and cloning.

