## Supplemental Figures and Tables for "A polarized nuclear position is required for correct division plane specification during maize stomatal development"

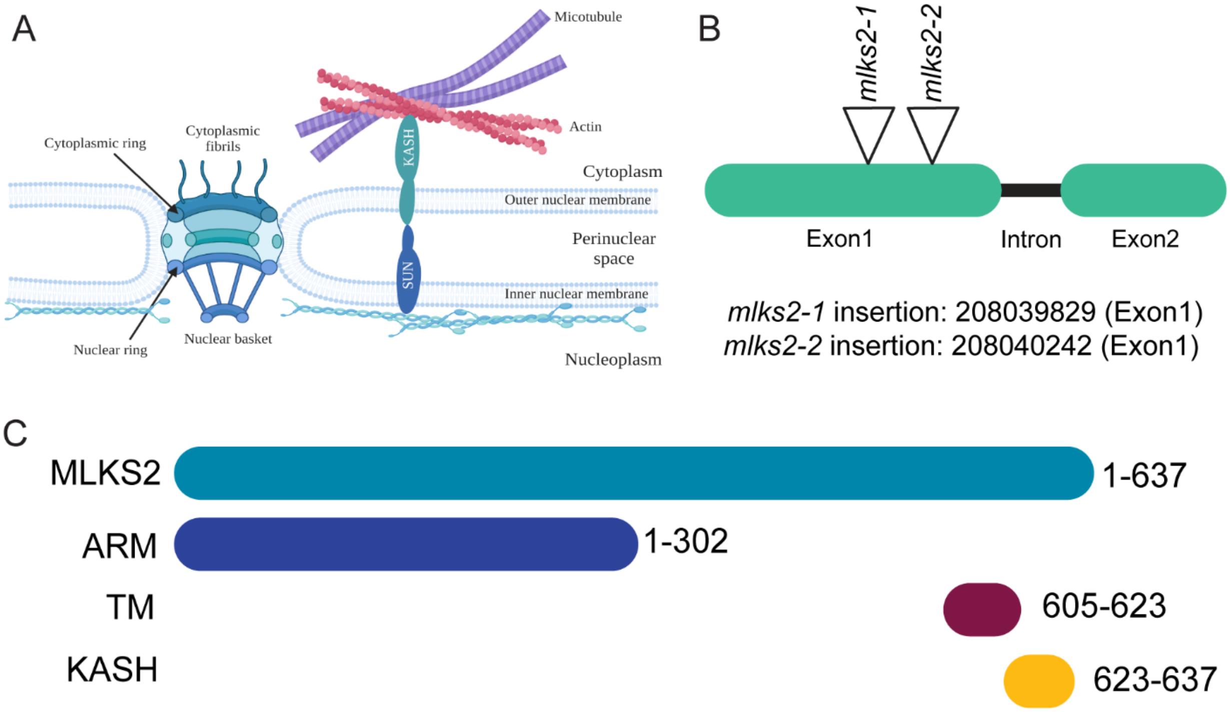


**Supplemental Figure S1. MLKS2 is an outer nuclear membrane protein and part of the LINC complex.** A, KASH-domain containing outer nuclear membrane protein interacts with SUN-domain containing inner nuclear membrane protein in the perinuclear space. KASH-domain containing outer nuclear membrane protein MLKS2 interacts with cytoskeleton. B, Position of two Mu insertion lines, *mlks2-1* and *mlks2-2*, of *Mlks2*. C, Domain organization of MLKS2 protein: Armadillo domain (1-302 aa), transmembrane domain (605-623), and KASH domain (623-637).

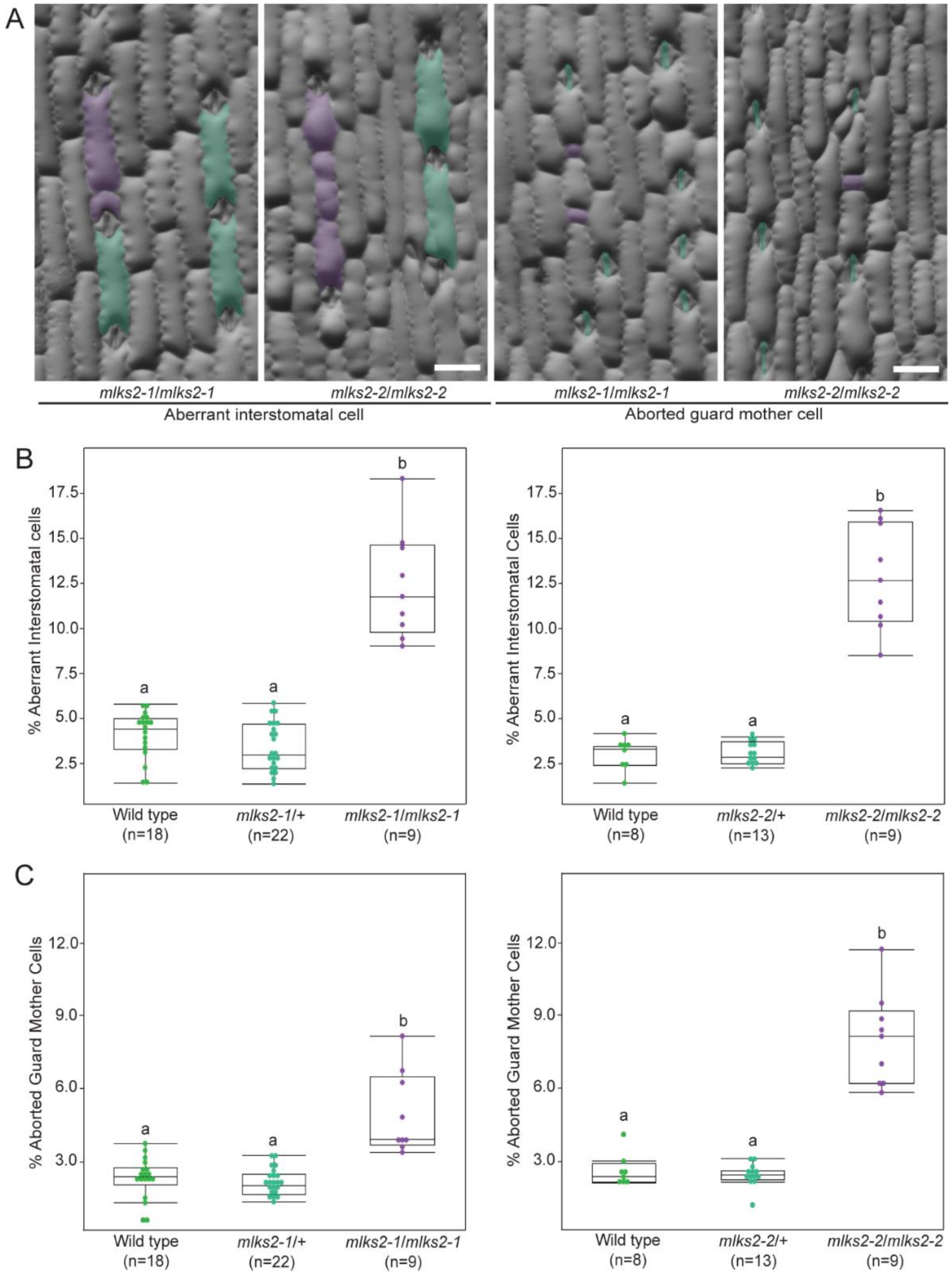


**Supplemental Figure S2. MLKS2 is required for proper interstomatal cell and guard cell division.** A, Interstomatal cell and aborted guard mother cell phenotypes of *mlks2-1* and *mlks2-2*. Cyan and magenta highlight indicate the normal and aberrant interstomatal cell or aborted guard mother cell, respectively. Scale bar = 0.1 mm. Same scale bar is applicable to all images. B, Quantification of aberrant interstomatal cell of *mlks2-1* and *mlks2-2*. C, Quantification of aborted guard mother cells of *mlks2-1* and *mlks2-2*. Number of plants (n) used per genotypes are mentioned. At least 200 cells are observed from each individual plant. Statistical test is performed based on Tukey's Honest Test. Groups labelled with the same letter are not statistically different from each other (alpha = 0.05).

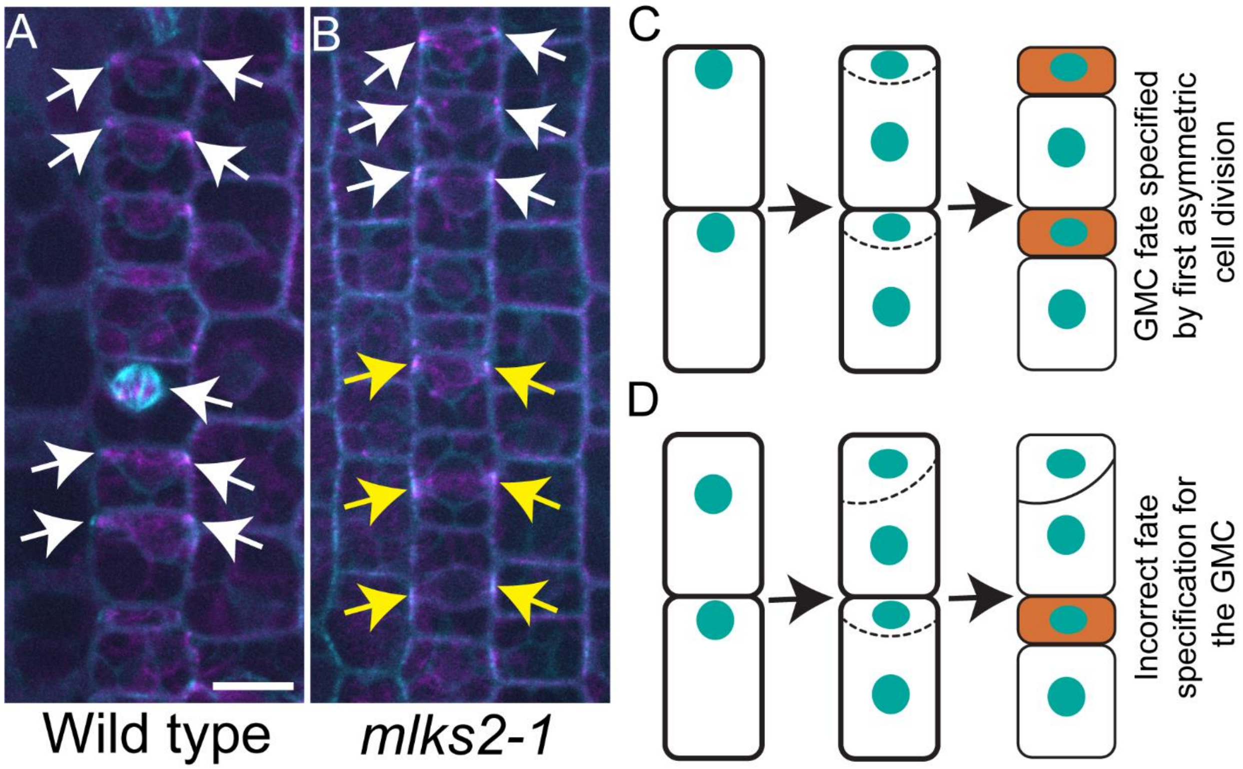


**Supplemental Figure S3. MLKS2 regulates the first asymmetric cell division.** First asymmetric cell division in wild type (A) and *mlks2-1* (B). Observation of PPB formation and nuclear position from cortex (magenta) and midplane (cyan), respectively, using CFP-TUB in wild type and *mlks2-1*. White and yellow arrow indicate the correct orientation and misorientation of first asymmetric cell division, respectively, in A and B. Scale bar = 10 μm. Same scale bar is applicable to all images. Guard mother cell (GMC) fate specification models based on first asymmetric cell division (C and D). C, Correct orientation of first asymmetric cell division results into the fate specification for GMC and interstomatal cell. D, Misorientation of first asymmetric cell division causes incorrect fate specification for GMC and interstomatal cell.

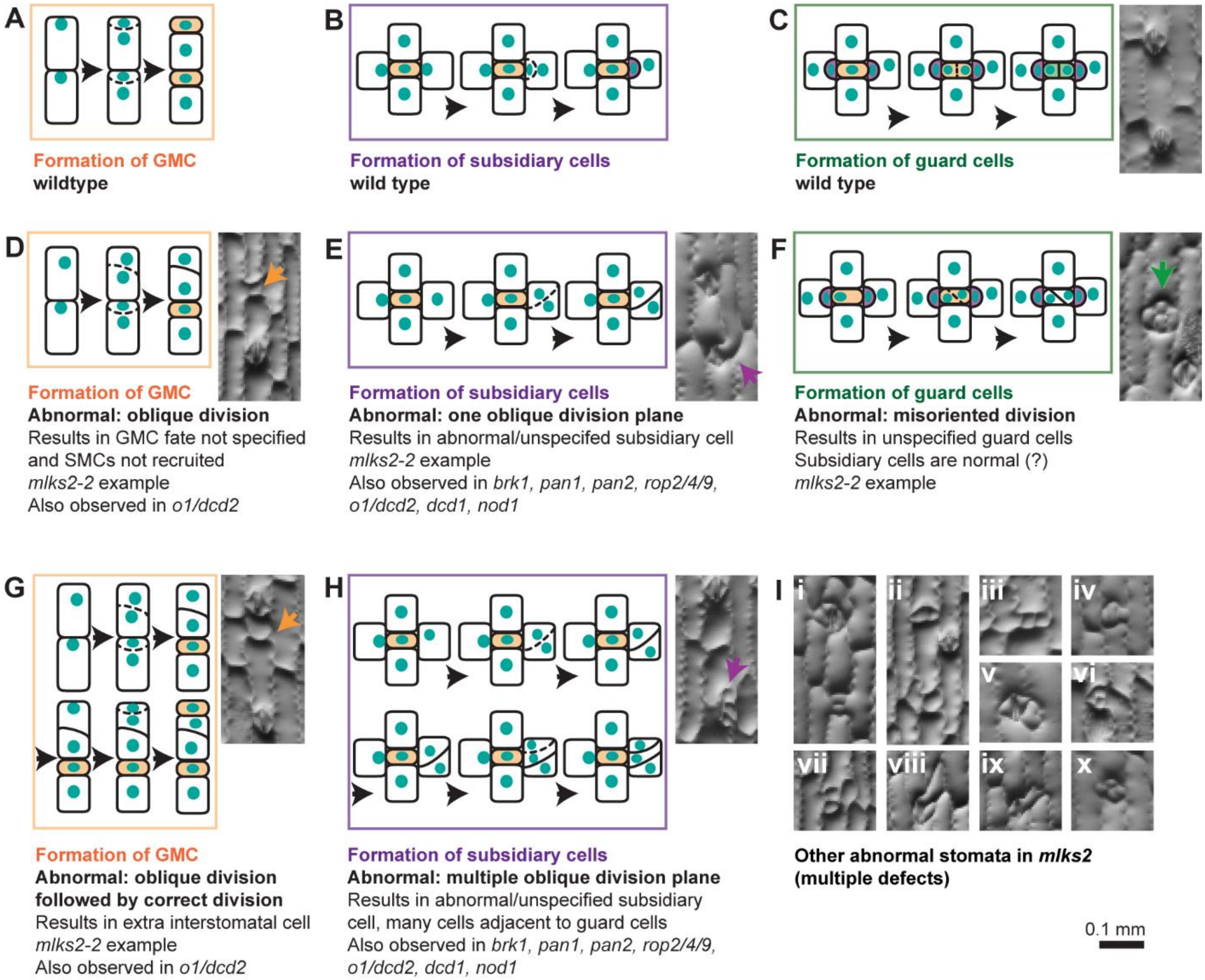


**Supplemental Figure S4. Different abnormal divisions result in different stomatal phenotypes in *mlks2*.** Progression of divisions that occur to produce (A-C) normal stomata or (D-H) abnormal stomata. (A) Asymmetric divisions that form the guard mother cell (GMC) and a sister interstomatal cell. Two divisions are shown in the cartoon. (B) Asymmetric divisions of the subsidiary mother cell (SMC) that produces a subsidiary cell and a pavement cell. (C) Symmetric, oriented division of the GMC forms two guard cells. Final panel shows a pair of normally formed stomata separated by a single interstomatal cell. (D, G) Abnormal division planes lead to a GMC that is not specified (C) or an extra interstomatal cell (G). (E, H) Abnormal division planes of the SMC lead to abnormal or unspecified subsidiary cell. In H, the SMC divides abnormally, and then divides again, leading to a cluster of abnormally shaped cells. The second division that occurs after an initial abnormal division may be abnormal, as in H. The second division can also be normal, resulting in a normal stomatal complex, and a small adjacent cell (See panel Iv). (F) Abnormal division plane of the GMC, resulting in unspecified guard cells. Note all cells have crenulations in the example photograph, suggesting all cells have a pavement cell fate. (I) Examples of irregularly shaped, misspecified, or improperly arranged cells observed in *mlks2*. Many architectures (such as iii, vi, iv and x) are unique to mlks2, and involve abnormal guard cell-generating divisions. A single complex may have more than one defect (e.g., iii and vi).

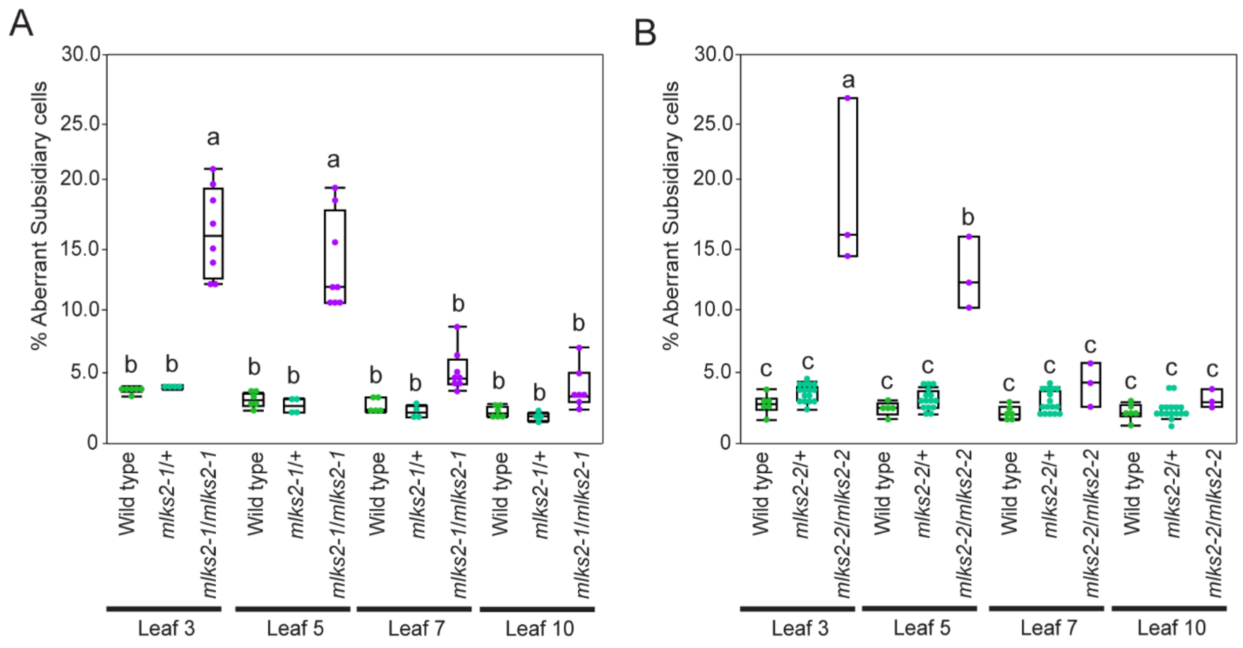


**Supplemental Figure S5. Aberrant subsidiary cell frequency of *mlks2-1* varies based on growth condition.** Quantification of aberrant subsidiary cells from leaf 3, 5, 7, and 10 of *mlks2-1* (A) and *mlks2-2* (B) grown in the summer field. At least 200 cells are observed from each individual plant. Statistical test is performed based on Tukey's Honest Test. Groups labelled with the same letter are not statistically different from each other (alpha = 0.05).

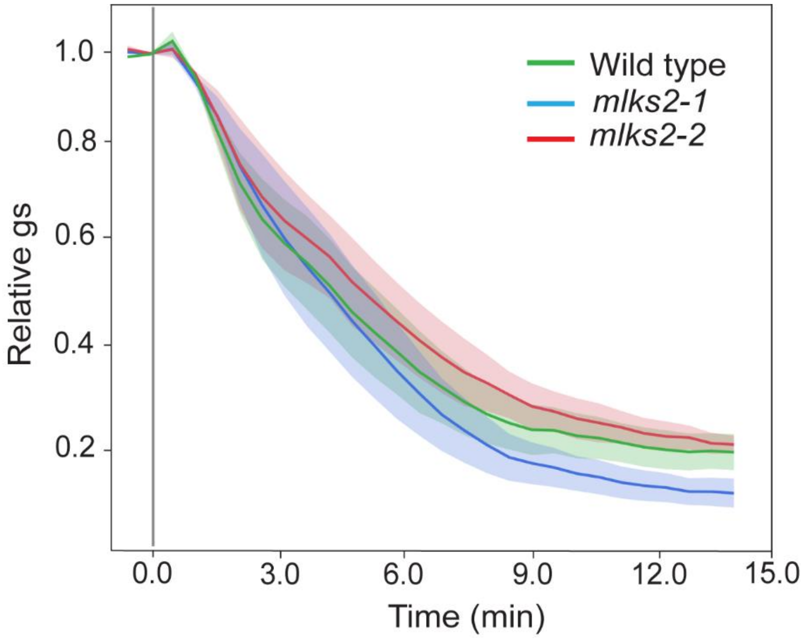


**Supplemental Figure S6. Stomatal conductance is not affected in *mlks2-1*.** Stomatal conductance of *mlks2-1* and *mlks2-2* were measured from leaf 3 using the CIRAS3 Portable Photo-Synthesis System. The data presented are the mean ± SE of at least 4 individual plants. Relative gs was computed for each individual measured plant by normalizing gs to the steady initial gs value observed.

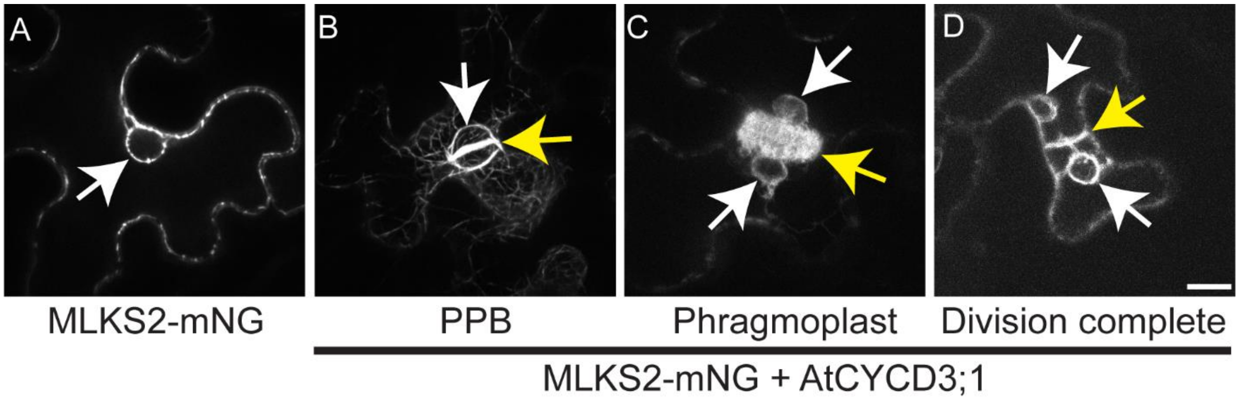


**Supplemental figure S7. MLKS2 localizes at the nuclear membrane and with mitotic apparatus.** A, Transient tobacco expression of MLKS2-mNG; where MLKS2 localizes at the nuclear membrane. After mitotic induction with AtCYCD3;1, MLKS2 also localizes at PPB (B), phragmoplast and new cell plate (C); and completes the mitosis (D). White arrow indicates nuclear membrane specific localization. Yellow arrow indicates PPB (B), phragmoplast and new cell plate (C); and completion of the mitosis (D). Scale bar = 20 μm. Same scale bar is applicable to all images.

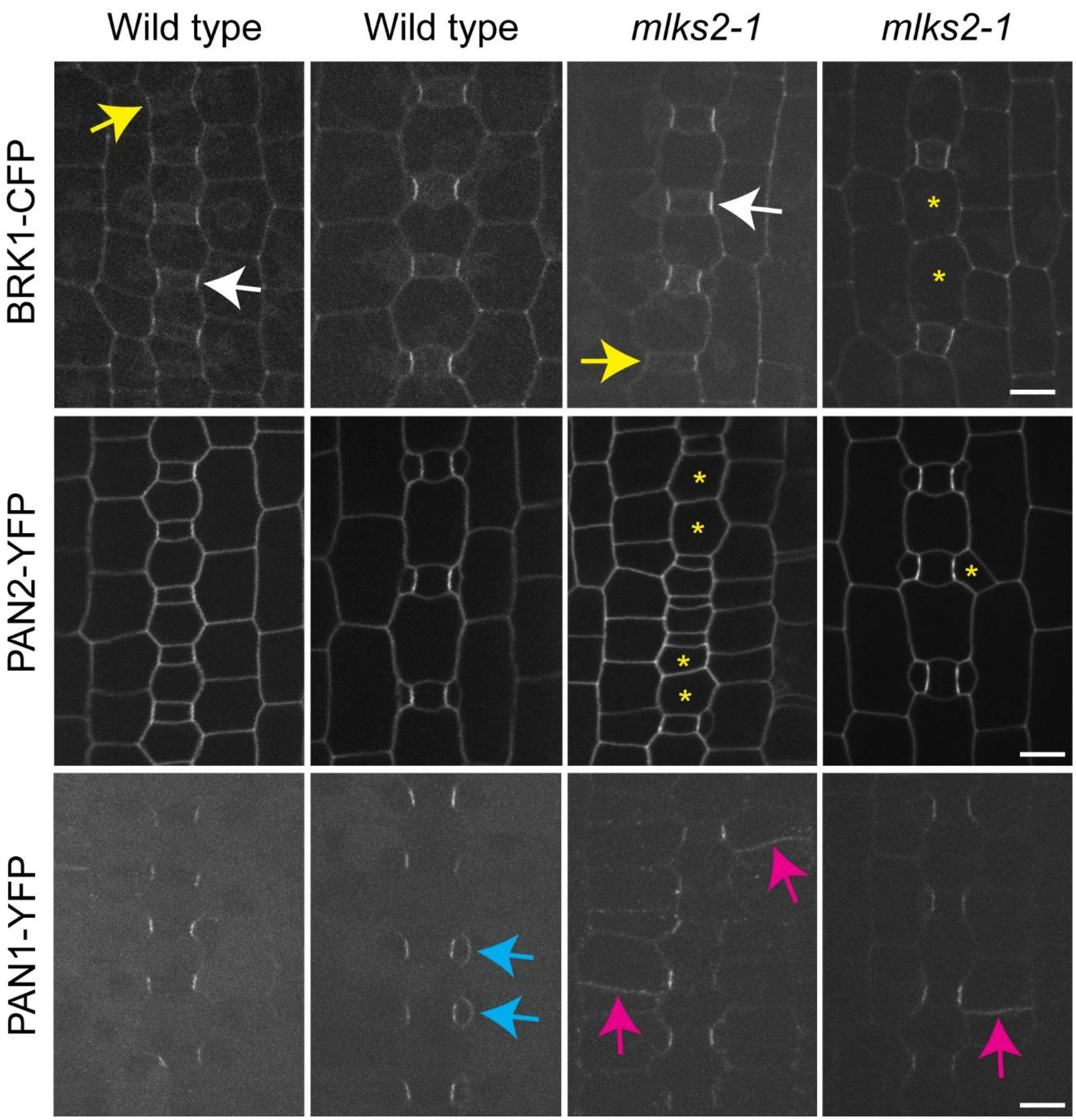


**Supplemental Figure S8. Earlier cell polarity markers remain the same in *mlks2-1*.** Localization of BRK1-CFP, PAN2-YFP, and PAN1-YFP in wild type and *mlks2-1* at two distinct developmental stages. For BRK1-CFP, white and yellow arrow indicate presence and absence of polar localization, respectively. Yellow asterisks in BRK1-CFP and PAN2-YFP highlight extra interstomatal cells or aberrant subsidiary cells in *mlks2-1*. For PAN1-YFP, cyan and magenta arrow indicate correct and incorrectly formed new cell plate in wild type and *mlks2-1*, respectively. Scale bar = 10 μm. Same scale bar is applicable to all images.

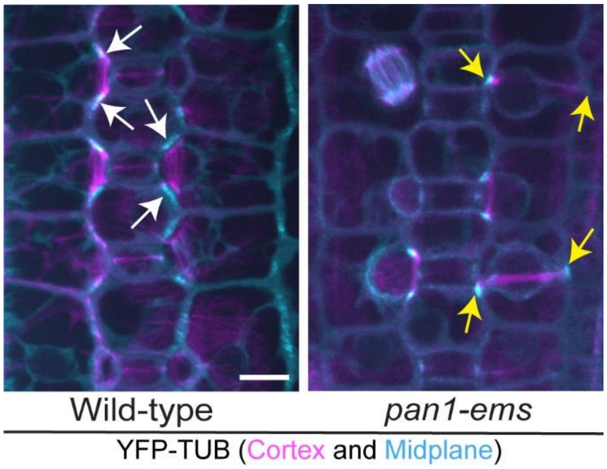


**Supplemental Figure S9. Transverse PPB formation in the *pan1-ems*.** Observation of PPB formation and nuclear position from cortex (magenta) and midplane (cyan), respectively, using YFP-TUB in wild type and *mlks2-1*. White and yellow arrow indicate perfectly placed and misplaced PPB, respectively. At least 3 plants from each genotype and ~100 cells from each plant were observed. Scale bar = 10 μm. Same scale bar is applicable to all images.

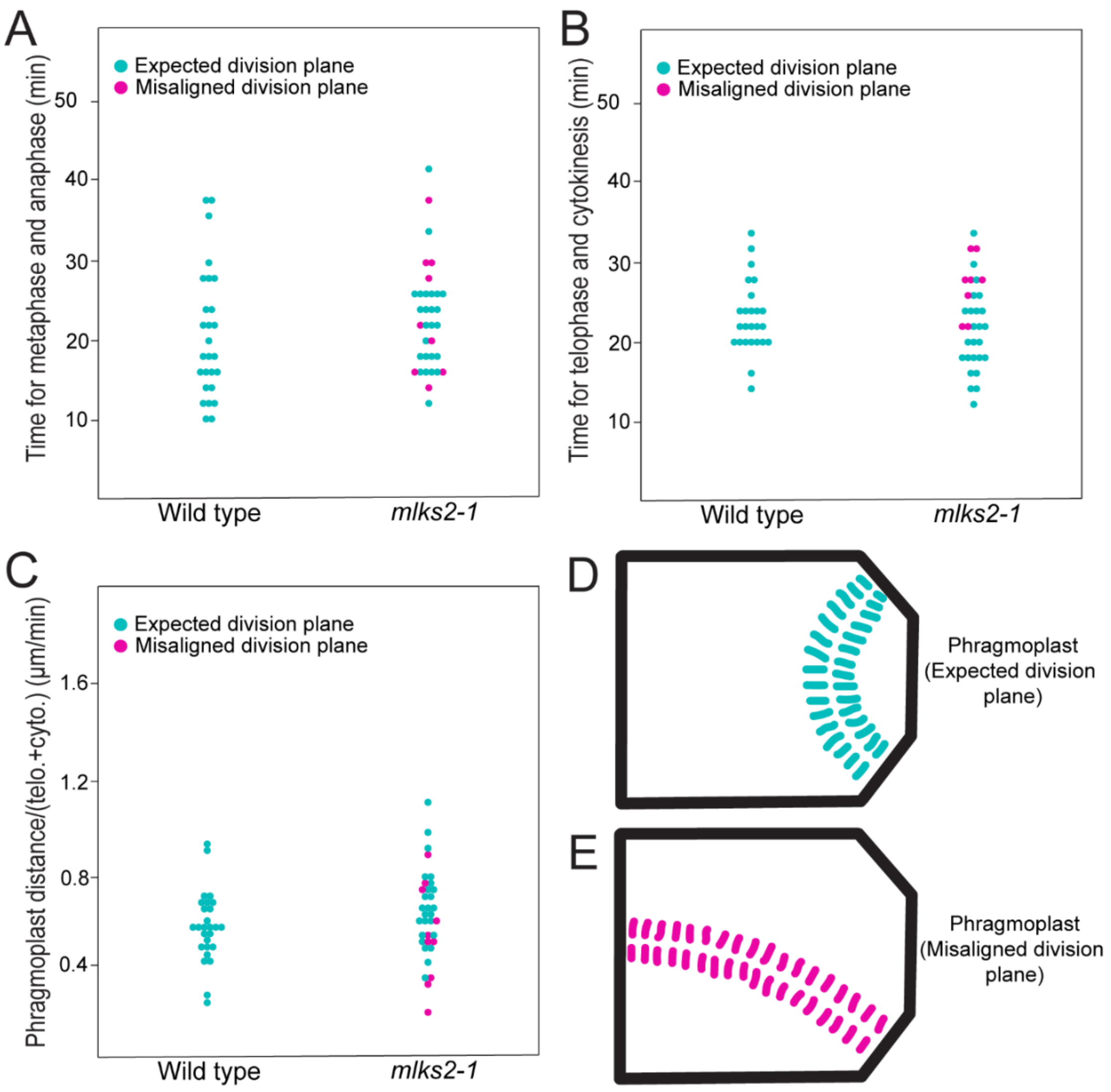


**Supplemental Figure S10. Mitosis timing is not altered in *mlks2-1*.** Duration of mitosis is divided into: metaphase + anaphase; telophase + cytokinesis. Duration of metaphase + anaphase (A) and telophase + cytokinesis (B) from the observed time lapse movies, related to Figure 6. C, Phragmoplast distance covered by each cell is measured as distance and divided by the time required for telophase + cytokinesis. Cyan and red dot indicate cells with expected division plane and misaligned division plane, respectively, from both wild type and *mlks2-1*. Student t-test has been performed for wild type and *mlks2-1*. P-values: 0.299 (A), 0.710 (B), and 0.222 (C). D, Cartoon shows the distance covered by phragmoplast for cells with expected division plane. E, Distance covered by phragmoplast for cells with misaligned division plane.

**Supplemental Table S1.** Primers used in this study for genotyping and cloning.

| Primer | Identifier | Sequence |
|---|---|---|
| MF03 | TIR6 | agagaagccaacgccawsgcctcyatttcgtc |
| MF533 | mlks21_gF1 | ggaggcgtacgcgaggtcca |
| MF534 | mlks21_gR1 | gaggctccggcatgggctct |
| MF535 | mlks22_gF2 | attcatccatcatgaacggcagcac |
| MF536 | mlks22_gR2 | ctgggactgggagtcttgtcccttg |
| AA106 | MLKS2_fwd_for_mNG | ggctac**cacctgc**agcg***ctgg***GGGGACAAGTTTGTACAAAAAAGC |
| AA107 | MLKS2_rev_for_mNG | ggctac**cacctgc**ttcc***TCCA***GCAGTGGGGACAAG |
| AA108 | mNeonGreen_fwd | ggctac**cacctgc**ttcc***tgga***gctgctgcaGCTGGTGCAATGGTGAGC |
| AA109 | mNeonGreen_rev | ggctac**cacctgc**tgcg***atgc***GGGGACCACTTTGTACAAG |
| AA98 | MLKS2_attB1_F | GGGGACAAGTTTGTACAAAAAAGCAGGCTATGGGCCGGAGCCTGAGCCCGC |
| AA47 | mNG_attB2_R | GGGGACCACTTTGTACAAGAAAGCTGGGTtcaCTTGTACAGCTCGTCCATGCC |

### Supplementary Movie Legends

**Supplemental Movie S1. MLKS2 localizes at the nuclear membrane and with phragmoplast and new cell plate**. Transient tobacco expression with pCaMV35S::MLKS2-mNG and pCaMV35S::AtCYCD3. Time lapse movie was captured after 48h of the transformation. The images were captured in every 2-minute interval.

**Supplemental Movies S2 – S10:** All Movies are played at 5 frames per second, with frames taken at 2-minute intervals. In all movies CFP-tubulin is green and YFP-Histone is magenta.

**Supplemental Movie S2. Normal asymmetric division of an SMC in a wild type sibling.**

**Supplemental Movie S3. Abnormal transverse division of an SMC in *mlks2-1*.**

**Supplemental Movie S4. Overview of the stomatal division zone in *mlks2-1*.** The video shows 3 stomatal rows and a variety of nuclear behaviors.

**Supplemental Movie S5. "TG" nuclear movement in a wild type SMC before PPB formation.** Normal asymmetric division of an SMC in a wild type sibling is shown in the top most SMC. In the bottom SMC, PPB formation has not yet occurred and the cell has an unpolarized nucleus with a stable position. The nucleus moves towards the GMC at the end of the video. Classified as "TG".

**Supplemental Movie S6. "AG" nuclear movement in a *mlks2-1* SMC before PPB formation**. The nucleus is initially polarized and touches the SMC wall closest to the GMC. The nucleus moves away and touches the side wall and eventually the wall distal to the GMC.

**Supplemental Movie S7. "TG/AG" nuclear movement in a *mlks2-1* SMC before PPB formation.** The nucleus is initially unpolarized, and then becomes offset, then polarized, and then moves away from the GMC to become unpolarized.

**Supplemental Movie S8. Stable polarized nuclear positioning in wild type SMCs after PPB formation.** Six SMCs are shown. The SMC in the bottom left is dividing, all other SMCs have PPBs and the nuclear position is stable. These nuclei are classified as "stay".

**Supplemental Movie S9. Unpolarized nuclei in SMCS after PPB formation.** Six SMCs are shown. The cell in the bottom right divides normally. All other SMCs have a PPB. Two nuclei are initially offset and move towards and away from the GMC.

**Supplemental Movie S10. "TG/AG" nuclear movement in a *mlks2-1* SMC after PPB formation.** An unpolarized nucleus is initially unpolarized and shows movements both towards and away from the GMC.
